# Xrn1 influence on gene transcription results from the combination of general effects on elongating RNA pol II and gene-specific chromatin configuration

**DOI:** 10.1101/2020.06.02.129171

**Authors:** Victoria Begley, Antonio Jordán-Pla, Xenia Peñate, Ana I. Garrido-Godino, Drice Challal, Abel Cuevas-Bermúdez, Adrià Mitjavila, Mara Barucco, Gabriel Gutiérrez, Abhyudai Singh, Paula Alepuz, Francisco Navarro, Domenico Libri, José E. Pérez-Ortín, Sebastián Chávez

## Abstract

mRNA homeostasis is favored by crosstalk between transcription and degradation machineries. Both the Ccr4-Not and the Xrn1-decaysome complexes have been described to influence transcription. While Ccr4-Not has been shown to directly stimulate transcription elongation, the information available on how Xrn1 influences transcription is scarce and contradictory. In this study we have addressed this issue by mapping RNA polymerase II (RNA pol II) at high resolution, using CRAC and BioGRO-seq techniques in *Saccharomyces cerevisiae*. We found significant effects of Xrn1 perturbation on RNA pol II profiles across the genome. RNA pol II profiles at 5’ exhibited significant alterations that were compatible with decreased elongation rates in the absence of Xrn1. Nucleosome mapping detected altered chromatin configuration in the gene bodies. We also detected accumulation of RNA pol II shortly upstream of polyadenylation sites by CRAC, although not by BioGRO-seq, suggesting higher frequency of backtracking before pre-mRNA cleavage. This phenomenon was particularly linked to genes with poorly positioned nucleosomes at this position. Accumulation of RNA pol II at 3’ was also detected in other mRNA decay mutants. According to these and other pieces of evidence, Xrn1 seems to influence transcription elongation at least in two ways: by directly favoring elongation rates and by a more general mechanism that connects mRNA decay to late elongation.

## Introduction

The dynamic equilibrium that keeps mRNA concentration is a balance between synthesis and degradation processes. In the classic paradigm, the ‘central dogma’ of molecular biology is a linear path in which the mRNA plays a role as an intermediary between genes and proteins. In the recent years our team has helped to demonstrate that there is a general control of the total cellular concentration of mRNA ([mRNA] homeostasis) in the yeast *Saccharomyces cerevisiae* and that it is the result of the crosstalk between transcription and degradation machineries (mRNA-crosstalk) (1). This involves both the mRNA imprinting by RNA binding proteins or base methylation, during its synthesis (2), as well as the feedback on the transcription machinery exercised by the decaysome (1). Other authors have demonstrated the existence of additional cross-talk mechanisms in this yeast dependent on the Ccr4-Not complex (3) or on the Nab2 mRNA binding protein (4). There are also evidences of the existence of transcription-degradation cross-talk in human cells for a subset of mRNAs through the Ccr4–Not and Xrn1 complexes (5–7) and during viral infections (8). In fact, for several of these complexes it has been demonstrated that there is also an extended connection to translation of the imprinted mRNAs. Both the Ccr4-Not (9) and the Xrn1 (decaysome) complexes (10) have a positive role on the translation of certain mRNAs that were, presumably, previously imprinted by them during transcription. This points to the existence of a transcription-translation cross-talk as well.

RNA pol II transcription is traditionally divided into three phases (11): i) the formation of the PIC (pre-initiation complex) that requires the successive binding of a plethora of factors over the basal promotor sequences; ii) the elongation phase after the escape of RNA pol II, and associated factors, from the promoter while synthesizing the mRNA; and iii) the termination phase that is composed by a polyadenylation/cleavage step of the primary transcript and a release of the downstream RNA pol II by the concerted action of pause sequences and the activity of the Rat1 (Xrn2) exonuclease (see 12, 13 for a review).

After initiation, some auxiliary factors remain bound to the basal promoter and some others travel with RNA pol II and a new set of factors that are required for fast and accurate elongation (14–16). For instance, lack of the elongation factor TFIIS provokes increased levels of backtracked RNA pol II (17) and specific defects in RP genes (18). The elongation of RNA pol II through chromatin is influenced by the nucleosomal organization and specific histone modifications of each gene and, at the same time, elongation affects both nucleosome positions and histone marks (19). There is a continuous cross-talk between the elongating complex and the chromatin template. RNA pol II changes its appearance because of the multiple post-translational modifications (phosphorylation and others) that the carboxyterminal domain (CTD) of the larger subunit (Rpb1) undergoes along the transcription cycle (20). The different phosphorylated forms of CTD make a code that is recognized by specific factors along the different elongation sub-phases (20). The last part of the elongation phase starts with the transcription of a sequence that acts as a signal for the cutting and polyadenylation machinery (21). The most important sequence element for transcriptional termination is the polyadenylation signal (pAS), which, in yeast, consists of three sequence elements: the AU-rich efficiency element (EE), the A-rich positioning element (PE) and the U-rich elements located upstream (UUE) or downstream (DUE) of the cleavage site (21, 22). The latter is defined by a pyrimidine followed by multiple adenosines Y(A)n (22). These elements are bound, respectively, by the cleavage factor 1B (CF1B, Hpr1), 1A (CF1A: Clp1, Pcf11, Rna14 & Rna15) and the cleavage and polyadenylation factor (CPF: 13 subunits). The RNA is then cut, whereupon the coding portion is polyadenylated and the 3’ product rapidly degraded (13, 21). The elongation complex undergoes a structural modification during these events. Some elongation factors leave the complex shortly before the pAS whereas other remain bound (14). This transition is reflected in the profile of active RNA pol II molecules in the close vicinity before and after the pAS (23).

Xrn1 is a very large (175 kDa, 1528 amino acids in *S. cerevisiae*) protein with pleiotropic function present in all eukaryotes. It was originally cloned as an enhancer of karyogamy defects (*KEM1*; (24)) and then isolated again as a DNA-strand transferase (*DST2*; (25)) together with *DST1* (26), a gene that was shown to encode TFIIS (27). Finally, its name was changed into Xrn1 (eXoRriboNuclease 1) because of its 5’-3’ exonuclease activity (28). The structure of the *Klyuveromyces lactis* and *Drosophila* Xrn1 were determined in 2011 (29, 30). They showed that the nucleolytic activity is on the N-terminal whereas the large C-terminal moiety is not directly involved in nuclease activity.

A fundamental question is how Xrn1-decaysome affects the transcription process. Comparison of the effects of *xrn1Δ, ccr4Δ* and other mRNA decay- and transcription-related mutants on *GAL1*, using computational modeling, indicates that Xrn1 and Ccr4 transcriptional functions take place in parallel rather than in a concerted manner (31). The Ccr4-Not complex has been shown to bind transcription initiation and elongation complexes, and to counteract RNA pol II backtracking (9, 32). The information available for the Xrn1-decaysome is much less abundant and in some cases, contradictory. In order to cast light on how Xrn1 contributes to transcription we have performed a series of genome-wide and single-gene experiments using various methods to map the distribution of RNA pol II under Xrn1-perturbed conditions. Our results indicate that the absence of Xrn1 affects transcription during early and late steps of elongation and we suggest that this is an important component of mRNA buffering.

## Materials and methods

### Strains

*S. cerevisiae* strains used in this study are listed in Table S1. Strains were grown at 30°C, with liquid cultures growing in an orbital shaking incubator at 180 rpm in YP medium (1% yeast extract, 2% peptone) or SC - TRP medium (0.17% yeast nitrogen bases (YNB), 0.5% ammonium sulphate and amino acids leucine, histidine, lysine and nitrogenous bases adenine and uracil) and containing either 2% glucose (YPD) or 2% galactose (YPGal) as a carbon source.

For transcriptional shut-off assays, cells were grown in YPGal until reaching an OD_600_ of 0.2-0.3. At this point, 4mM of 4-thiouracil (Sigma) or DMSO was added to the media and left to grow for an additional 2 hours until reaching OD_600_ of approximately 0.5. Then, a sample was taken, and immediately after, glucose 4% was added to the culture. Samples were taken at the indicated times after glucose addition.

For obtaining CRAC samples, strains were grown in SC –TRP until an OD_600_ of 0.5-0.6. In parallel *Schizosaccharomyces pombe* was also grown to an OD_600_ of 0.5-0.6 in minimal media (EMM 0.5X), in order to add 1% to the culture before cross-linking as a spike-in. To the XRN1-AID strain 0.2mM of auxin (3-Indoleacetic acid; Sigma-Aldrich) was added to the media 30 and 60 minutes before harvesting. To the XRN1-AA strain 1 µg/ml of rapamycin (Sigma) was added to the media 60 minutes before harvesting.

### RNA extraction, RT-qPCR and mRNA half-life calculation

Cells were grown to log phase and 10 ml was harvested by centrifuging for 2 minutes at 4000 rpm and flash freezing the cells in liquid nitrogen. Total RNA was purified using hot phenol–chloroform method as described previously (33) and reverse-transcribed to cDNA using M-MLV Reverse Transcriptase (Promega). A relative real-time quantitative PCR was then carried out for all the genes studied (Table S2) against *SCR1* using SYBR Green Premix Ex Taq (Takara) in a Light Cycler 480 II (Roche).

To determine the mRNA half-lives, we used a transcription shut-off assay, collecting samples at 5, 20 and 50 minutes after glucose addition. mRNA levels were then determined with RT-qPCR and half-lives were estimated by calculating the time it takes for half of the initial amount of RNA to be degraded. Primers used are listed in Table S2.

### Chromatin immunoprecipitation (ChIP)

Yeast strains were grown to exponential phase and 50 ml were taken for each sample. The ChIP experiments were performed as previously described (34), except that anti-Rpb3 (ab81859; Abcam), anti-Ser2P (ab5095; Abcam), Anti-Ser5P (ab5131; Abcam), anti-Tyr1P (3D12; Merck), anti-H3K4me3 (ab8580; Abcam) or anti-H3K36me3 (ab9050; Abcam) antibodies and Sheep anti-Mouse IgG Dynabeads (Invitrogen) were used. Genes were analysed by quantitative real-time PCR in triplicate using primers listed in Table S2.

The ChIP-seq data was generated in a separate study (31).

### CRAC

RNA pol II CRAC was generated and analysed as in (35). Briefly, cells grown until exponential phase were crosslinked at UV 254nm for 50s and harvested by centrifugation. The cells were then resuspended in TN150 buffer (50mM Tris pH 7.8, 150mM NaCl, 0.1% NP-40 and 5mM β-mercaptoethanol) supplemented with protease inhibitors (complete, Mini, EDTA-free Protease Inhibitor Cocktail). This suspension was flash frozen in droplets and cells were mechanically broken using the Mixer Mill MM 400. The resulting cell “powder” was thawed and treated for one hour at 25 °C with DNase I (165 U/g of cells). The cells were centrifuged and subsequent purifications steps were performed. The purified complexes were eluted with TEV protease and treated with an RNase cocktail (RNace-IT, Agilent) to reduce the size of the nascent RNA. The adaptors were modified in order to sequence RNA molecules from the 3’-end. The RNA was recovered after proteinase K treatment by phenol-chloroform extraction and reverse transcribed using specific primers. The resulting cDNA was amplified with multiple PCR reactions in a final volume of 25 μl using the following conditions: 0.4 μM of each primer, 0.2mM dNTP, 2.5 U LA Taq DNA polymerase from Takara, 1x LA PCR Buffer II and 2 μl of complementary DNA per reaction with the programme: 2’ at 95 °C, (30’’ at 95 °C, 45’’ at 58 °C, 1’ at 72 °C) × 13 cycles, 5’ at 72 °C. All the PCR reactions were pooled together and treated with 200 U of Exonuclease I (NEB) per ml of PCR reaction for 1 h at 37 °C. After Exonuclease I inactivation for 20’ at 80 °C, DNA was purified with PCR clean up columns (NucleoSpin Gel and PCR Clean-up, Macherey-Nagel) and sequenced using Illumina technology.

### BioGRO-seq

The Bio-GRO protocol was essentially as described in (36) for tiling array analysis, but modified for high-throughput sequencing. For each sample, 100-ml cultures were grown to OD_600_ = 0.55. Cells were collected by centrifugation and frozen in liquid nitrogen. Frozen pellets were transferred immediately to -20 °C. After at least 3 h, cells were thawed on ice and permeabilized with 10 ml of 0.5 % sarkosyl solution. RNase trimming was achieved by incubating cells with 32 µl of RNase A (10 mg/ml) dissolved in 3.2 ml of 0.5% sarkosyl solution under agitation conditions for 10 min at 30 °C. In order to remove RNase, cells were washed three times with 50 ml of 0.5 % sarkosyl and transferred to an eppendorf tube. Cells were resuspended in 120 µl of water plus 5 µl of RNase OUT. The run-on reaction was performed by adding 120 µl of transcription buffer 2.5X (50mM Tris–HCl, pH 7.7, 50mM KCl, 80mM MgCl_2_), 16 µl of ACG mix (ATP, CTP, GTP, 10 mM each), 6 μL of 0.1M DTT and 20.25μL of Bio-11-UTP (10 mM, Metkinen) and incubating the mixture at 30 °C for 5 min. The reaction was stopped with 1 mL of cold water. After run-on labelling, cells were kept on ice for 5 min and harvested by centrifugation. RNA extraction was done using the ‘MasterPure Yeast RNA Purification Kit’ (Epicentre) following the manufacturer’s instructions. Once extracted, contaminating genomic DNA was removed by incubating with 2 µl of RNase-free DNase I (Roche) for 30 min at 37 °C. Purified RNA was resuspended in 32 µl of water and was spectrophotometrically quantified. Labelled RNA was subjected to 1 round of affinity purification with streptavidin paramagnetic beads (Dynabeads M-280 Streptavidin, Invitrogen). Purified labelled RNA was quantified with Qubit (Invitrogen) and 5’-OH ends were repaired with PNK prior to library preparation with the NEBNext^®^ Small RNA Library Prep Set for Illumina^®^ (Multiplex Compatible).

### Nucleosome mapping with MNase

The MNase digestion was performed as described in (37). Briefly, 500 ml of exponentially grown yeast culture was cross-linked with formaldehyde. Cells were washed and resuspended in Buffer Z2. Zymolyase was added and cells were incubated for 30 min at 30 °C to obtain spheroplasts. Spheroplasts were incubated with different amounts of micrococcal nuclease (from 500 to 10 mU). Protein was degraded by Proteinase K and DNA was obtained by phenol-chloroform-isoamyl extraction and ethanol precipitation. Digested DNA was resolved on an agarose gel and the band corresponding to mononucleosomes was purified. Naked DNA was obtained as before but MNase was added after DNA extraction instead of before, and a band with similar digestion as the mononucleosomes was purified.

MNase-digested mononucleosomal fragments of DNA resulting from wild-type and *xrn1Δ* strains and the corresponding nucleosome-sized fragments of naked DNA were sent to the genomic facility of Centre of Genomic Regulation (CRG) in Barcelona to be sequenced. These DNA fragments were sequenced using paired-end technology from Illumina (HiSeq Sequencing v4 Chemistry with read length 50).

### High resolution polyA-site mapping

We used unpublished *xrn1Δ* polyadenylation site mapping datasets generated by the internal version of the 3’T-fill method (37) to map the polyA sites at high resolution. The accession number for those data is GSE158548. As a *wt* reference, we downloaded and processed raw sequencing files from GSE40110.

### Bioinformatics

CRAC samples were demultiplexed using the pyBarcodeFilter script from the pyCRAC utility suite. Subsequently, the 3’ adaptor is clipped with Cutadapt and the resulting insert is quality trimmed from the 3’-end using Trimmomatic rolling mean clipping (window size =5, minimum quality =25). At this stage, the pyCRAC script pyFastqDuplicateRemover is used to collapse PCR duplicates and ensure each insert is represented only once. Each unique insert in our library is associated with a six nucleotides random tag within the 5’ adaptor. The resulting sequences are reverse complemented with Fastx_reverse_- complement (part of the fastx toolkit), and mapped to the R64 genome (sgd) with bowtie2 (-N 1 –f) (38). Read counts were normalized relative to reads derived from an *S. pombe* spike-in. The *S. pombe* spike cells contain a non-relevant protein tagged with the same HTP tag that was co-purified with the *S. cerevisiae* sample. We then performed a metagene analysis using a package available for Python, ngsplot. We obtained the scatterplots, boxplots and density plots by counting the number of reads within the ORF of a gene using HTSeq, and then plotting this data in R. IGV (39) was used to directly compare reads at different positions in the genome.

AfterQC v0.9.6 (40) was used with default parameters to do the quality control, filtering and trimming of the paired-end sequencing data obtained from MNase. The cleaned sequence data was aligned to the S. cerevisiae (R64) reference genome with bowtie2-2.3.3.1 (38). Once mapped, the resulting SAM files were analysed using DANPOS v2.2.2 (41) in order to determine the nucleosome localization, occupancy and fuzziness across the whole genome in all the analysed strains. DANPOS pools the replica of the same condition and can use naked data as a background correction.

For BioGRO-seq samples, Trimmomatic (42) was used to clip sequencing adapters, trim reads with low quality scores and filter reads shorter than 20 bases. High quality reads were then aligned to the S. cerevisiae R64 reference genome with TopHat2. Normalised average density plots around genomic features were generated with the ngs.plot software (43). The RNA-seq mode and the statistical robustness parameter, which filters out 0.5% of genes with the most extreme expression values, were applied to all calculations.

For high-resolution polyA-site mapping, we used Trimmomatic to remove sequencing adapters, 5’-end barcodes, and 3’-end bases that fell below a minimum quality of 3. In order to more precisely remove polyA stretches, we used PRINSEQ (44) with the following parameters: min_gc 20, min_len 20, trim_tail_left 8, trim_tail_right 8, out_format 3. The remaining high quality reads were mapped to the sacCer3 (R64) reference genome with Bowtie2, with default parameters. Strand-specific bedgraph files were generated with the bamCoverage function from Deeptools (45), with the following parameters: bin size 1, normalization rpkm, scaling factor 1, ignore missing data false, ignore duplicates false, minimum mapping quality 1, offset -1. The last parameter was used to collapse the coverage track to the last 3’-end base of each read.

### Modelling of elongation rates

To estimate the RNA polymerase II elongation rates along the genomic region, we divide the region into spatial bins *i* ∈ {0,1,2, …} where *i = 0* corresponds to the Transcription Start Site (TSS). The RNA polymerase II hops from bin *i* to *i+1* with rate *k*_*i*_ that can be interpreted as the elongation rate at that genomic region, and we assume there is no polymerase drop off. Let *R*_*i*_ denotes the probability of RNA polymerase II occurrence in the spatial bin *i* where *i* ∈ {0,1,2, …}. Then, the net inflow rate of polymerases into bin *i+1* is *k*_*i*_*R*_*i*_(1−*R*_*i*+1_)which is proportional to the product of the probability that site *i* is occupied, and site *i+1* is empty. Similarly, the net outflow rate of RNA polymerases from bin *i+1* is *k*_*i*+1_ *R*_*i*+1_ (1−*R*_*i*+2_). If *k*_*TSS*_ is the arrival rate of RNA polymerase II to the TSS, one can write the following iterative equations based on the same inflow and outflow of polymerases at each location

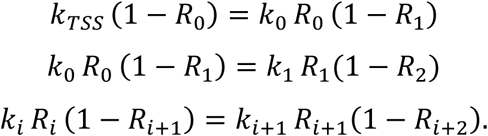

Further assuming that *R*_*i*_≪1 the above equations simplify as

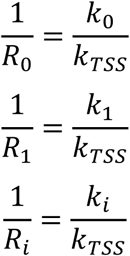

and the elongation rate *k*_*i*_ normalized with *k*_0_ is simply equal to the ratio *R*_0_/*R*_*i*_. Assuming *R*_*i*_ is proportional to the mean number of reads, we plot *R*_0_/*R*_*i*_in Fig. 1D as a function of *i*.

**Figure 1.**
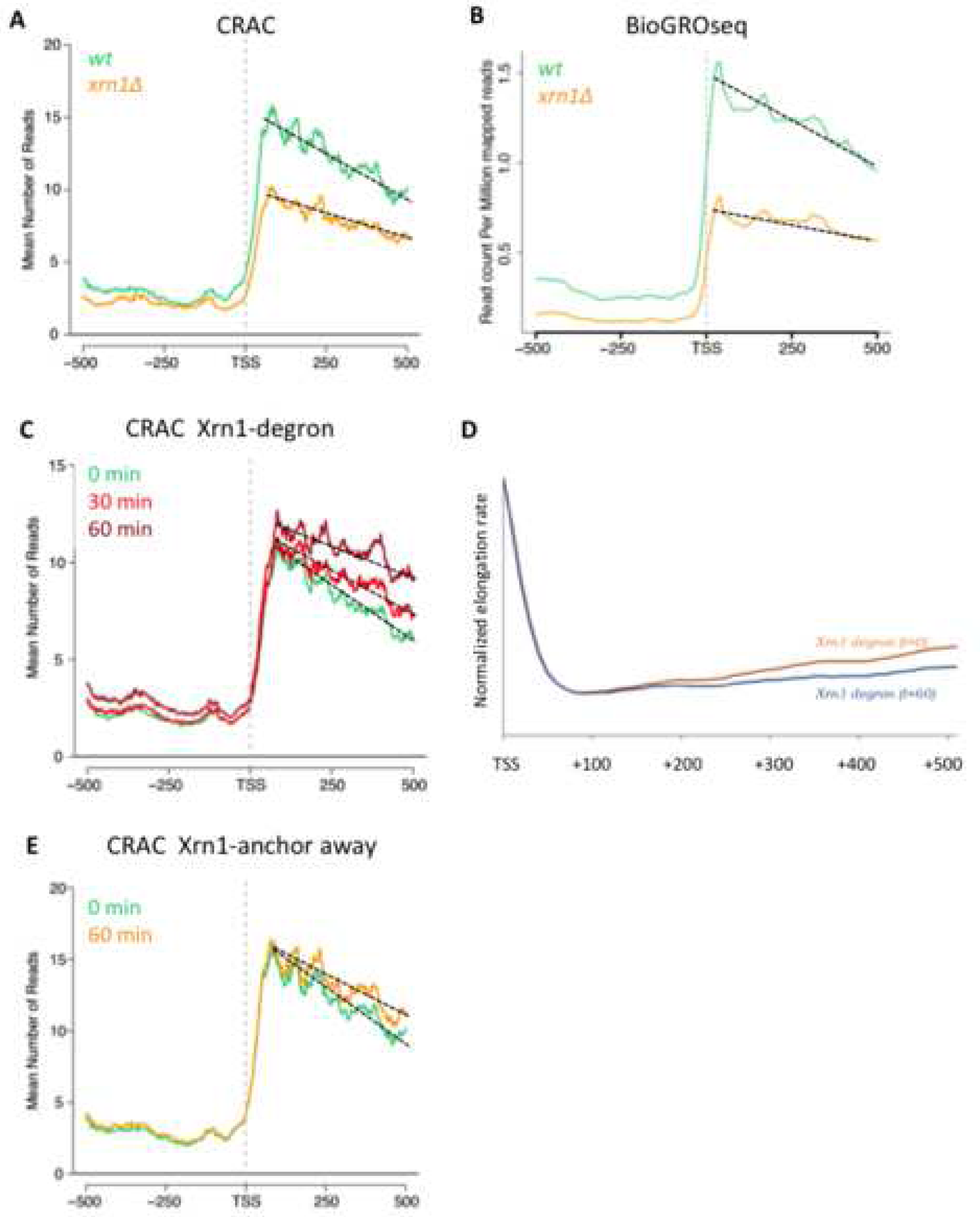
Xrn1 perturbation alters RNA pol II profiles downstream of the TSS. A) A metagene plot of the CRAC data for all canonical TATA-containing genes aligned by their transcription start site (TSS) shows that there is a global reduction in RNA bound to total RNA pol II and a decrease in the slope in a *xrn1Δ* strain (in orange) compared to the wild type (in green). Similar plots for all genes and for some specific gene groups are shown in Fig S3. B) A similar metagen plot of BioGRO-seq data for all TATA genes shows the same effect as in A for the nascent mRNA produced by active RNA pol II. C) The fast depletion of Xrn1 using the auxin-degron system shows a progressive change in the slope of the CRAC data of all TATA genes but with a small increase in the global number of reads. D) Estimation of elongation rates from the CRAC data after fast depletion shown in C, after solving iterative equations based on the same inflow and outflow of polymerases at each location (see Methods). E) The fast depletion of Xrn1 from the nucleus using the anchor-away protocol (orange) also shows a change in the slope compared to the wild type (green). A similar plot for all genes is shown in Fig S5B.

### Western blot

Laemmli-boiled crude extracts were run on a SDS-polyacrylamide gel and transferred to nylon membranes (Hybond-ECL). After blocking with Tris-buffered saline containing 0.1% Tween 20 and 5% milk, the following primary antibodies were used: anti-FLAG M2 (Sigma-Aldrich), anti-GFP (Roche) and anti-L1. Finally, the membrane was incubated with peroxidase-conjugated goat anti-mouse IgG and proteins were detected using a chemiluminescence detection kit (Super-Signal West Pico, Pierce) and a ChemiDoc™ MP system (Bio-Rad).

### Sucrose gradient centrifugation

For polysome preparations 200 ml cultures were grown in YPD and harvested at an OD_600_ between 0.5 and 1. Cycloheximide was added to a final concentration of 0.1 mg/ml before harvesting for 10 minutes. Cell extracts were obtained by adding lysis buffer (10 mM Tris-HCl (pH 7.5), 100 mM NaCl, 30 mM and 0.2 mg/ml of heparin) and glass beads, and vortexing for 8 minutes at 4 °C. The supernatant was recovered after centrifugation and flash frozen. Thawed extracts (10 absorption units at 260 nm) were layered onto 11.2 ml of 7 to 50% linear sucrose gradients that were prepared in 50 mM Tris-acetate (pH 7.5), 50 mM NH4Cl, 12 mM MgCl2, 1 mM dithiothreitol. The gradients were centrifuged at 39,000 rpm in a Beckman SW41 rotor at 4°C for 2 h 45 min. Gradient analysis was performed with an ISCO UV-6 gradient collector and continuously monitored at 254 nm. Fractions were collected every 30 s for 9 min (18 in total).

For western analysis, proteins (0.5 ml of each fraction) were precipitated by addition of trichloroacetic acid to a final concentration of 10%, followed by incubation on ice for 10 min. Proteins were pelleted by centrifugation for 10 min at 4°C. Pellets were washed twice with 1 ml of ice-cold acetone and finally resuspended in 20 µl of 2X Laemmli buffer. Every two fractions were mixed together and aliquots of 20 µl were loaded onto 10% SDS-polyacrylamide gels and analyzed by western blot analysis.

## Results

### CRAC experiments confirm that lack of Xrn1 decreases transcription rates

Conflicting results on the effect of the *xrn1Δ* mutation on transcription rates have been described, based on different experimental approaches (Fig S1A-B). We have previously shown that genomic run-on (GRO) analyses indicate a global negative effect of *xrn1Δ* on transcription rates across the genome (1). In contrast, results obtained after RNA metabolic labelling (cDTA) with 4-thiouracil (4-tU) were interpreted to support a positive effect of *xrn1Δ* on transcription rates (46).

To address this apparent contradiction, we adopted a third experimental approach, using RNA pol II CRAC to map the distribution of elongating RNA pol II molecules at nucleotide resolution. For a quantitative analysis of the effect of the *xrn1Δ* mutation we normalized our data to a spike in obtained from the addition to each sample of an equivalent amount of *S*.*pombe* cells expressing a tagged protein that could be co-purified with yeast RNA Pol II (47). This analysis confirmed a very significant negative effect of *xrn1Δ* on transcription rates (Fig S1C-E). The similarity between CRAC and GRO data extended to the dependency of the *xrn1Δ* effect on transcription levels. Both in GRO and CRAC experiments, the negative effect caused by the mutation was slightly stronger for highly transcribed genes (Fig S1F).

We considered the possibility that the differences with the metabolic labelling study of Sun et al (46) might be due to an effect of 4-tU on mRNA stability. Indeed, although in that study 4-tU was used to estimate transcription levels, a significant fraction of 4-tU containing RNAs after the 6 minutes labelling step might be post-transcriptional and subject to degradation. We set out to assess whether degradation of these 4tU labelled RNAs is delayed in *xrn1Δ* cell, which could lead to an overestimation of transcription levels in these cells.

To this end, we measured the stability of RNAs derived from genes of the galactose regulon with a glucose-shutdown approach. As shown in Figure S2A, the half-life of 4-tU containing RNAs is significantly increased in *xrn1Δ* cells. This suggests that 4-tU inhibits, *in vivo*, the exosome-dependent 3’-5’ degradation pathway that takes over mRNA degradation in cells defective for the 5’-3’, Xrn1-dependent pathway. Consistent with this notion, levels of ncRNAs that are degraded by the exosome increased in wild-type cells treated with 4-tU (Fig S2C). These results strongly suggest that the increase in 4tU-labelled RNAs levels, previously observed in the *xrn1Δ* mutant by Sun et al (46) might be due to the stabilization of these RNAs rather than to increased transcription rates in *xrn1Δ* cells.

### Xrn1 regulates transcription elongation at the 5’ of genes

In addition to an overall effect on the number of transcribing RNA pol II molecules, lack of Xrn1 produced significantly altered RNA pol II profiles across the genome. Metagene analyses showed an increase in the density of elongating RNA pol II molecules mapped in the wild type shortly downstream of the transcription start site (TSS) and a progressive decline afterwards (Fig 1A). This decline was less pronounced in *xrn1Δ*, which can be appreciated in the difference in slopes of the two strains (50% reduction of the wild-type slope in *xrn1Δ*). The difference in slope was particularly clear in highly transcribed genes and those containing canonical TATA elements, but was also easily detected in other categories of genes (Fig 1A, Fig S3).

Different RNA pol II profiles between wild-type and *xrn1Δ* 5’ metagenes were also detected after performing a high-resolution genomic run-on analysis (BioGROseq) (Fig 1B). Fast depletion of Xrn1, using an AID degron approach (Fig S4), also provoked a change in the CRAC RNA pol II profile in the 5’ end metagenes, exhibiting a decrease in the slope which is visible after just 30 min depletion (Fig 1C).

These results support an influence of Xrn1 on transcription elongation, as previously proposed (1, 31, 48). Changes in the slope of RNA pol II profiles has been usually interpreted as the consequence of alterations in RNA pol II processivity. According to this, the less negative slopes in Xrn1-perturbed strains would indicate increased processivity and would involve a negative influence of Xrn1 on this aspect of transcription elongation. Alternatively, the different slope detected in Xrn1-perturbed cells may indicate alterations in elongation rates, as we have already found for *xrn1Δ* in kinetic experiments with a galactose-controlled long gene (31). In order to test this last possibility, we decided to model our Xrn1-degron CRAC data, following an iterative equation approach. As shown in Fig 1D, in both conditions (depleted and non-depleted) estimated elongation rates dropped at the same position from TSS (+80 bp). From there on, our model estimated a progressive increase in the elongation rate for the non-depleted condition. In contrast, this acceleration was almost absent in the Xrn1-depleted condition (Fig 1D).

In the case of Xrn1-depleted data, there was no overall decrease in the number of reads as in *xrn1Δ* (compare Fig 1A to 1C). This is compatible with a scenario where *xrn1Δ* cells exhibit both elongation and initiation defects, the latter being responsible for the general decrease in the number of transcribing RNA pol II molecules. In contrast, Xrn1 depletion by the AID degron system would provoke just the elongation defect, producing no decrease but a slight accumulation of RNA pol II molecules on the gene body, likely as a consequence of its reduced elongation rate. According to this view, the initiation defect detected in *xrn1Δ* might be related, at least partially, to the downregulation of RNA pol II-dependent transcription caused by slow growth (36,48). RNA pol II CRAC analysis was also performed after depleting the nuclear pool of Xrn1, using an anchor-away approach (Fig S5). After 1 h incubation with rapamycin, we found a slight decrease in the slope of the RNA pol II profile at 5’ (Fig 1E), similar to the one produced by total Xrn1 depletion (compare Fig 1C to 1E, and S3A to S5B). This modest effect detected might suggest that the functional influence of Xrn1 on transcription elongation depends on its presence in the nucleus. However, we should be very cautious when interpreting these results, since we did not find any change in the slope when longer treatments were performed (2 and 5 hours in the presence of rapamycin) and, therefore, we cannot rule out that nuclear Xrn1 depletion was partial and transient.

### Lack of Xrn1 produced changes in nucleosome density on gene bodies

We have seen that the change in the slope of CRAC profiles at 5’ metagenes produced by the lack of Xrn1 indicates general alteration of transcription elongation that are compatible with either increased RNA pol II processivity or decreased elongation rate. Any of these two possibilities would involve increased competition for occupancy between elongating RNA pol II molecules and nucleosomes along gene bodies. Slower elongation would involve higher dwell time for elongating RNA pol II molecules and, as a consequence of it, increased competition with nucleosomes. Increased processivity would also predict a higher number of elongating RNA pol II molecules competing with nucleosomes in distal positions.

In order to test this prediction we generated nucleosomal maps of cells lacking Xrn1, using an MNase-seq approach optimized for detecting elongation-related phenomena (49). These maps showed increased occupancy of the two nucleosomes flanking the nucleosome-depleted region (NDR) just upstream of the TSS in *xrn1Δ* (Fig 2A). This may be indicative of a lower frequency of PIC assembly, as a consequence of the competition between PIC assembly and nucleosome +1 (50), and is consistent with our hypothesis of overall decrease in transcription initiation in *xrn1Δ*. Alternatively, this might also be consistent with less pausing in the early stages of elongation.

**Figure 2.**
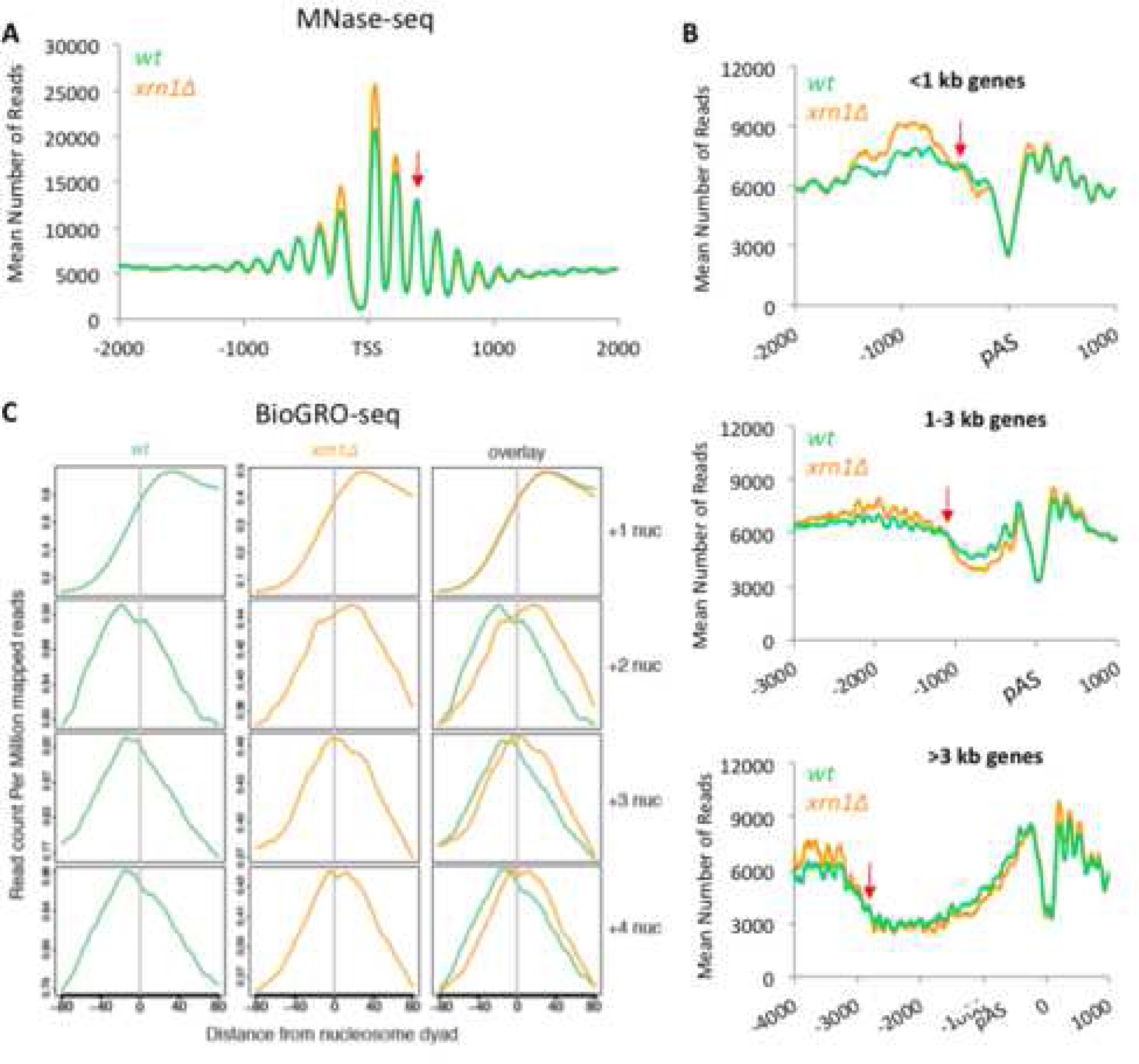
Lack of Xrn1 changes the chromatin landscape and the distribution of active RNA pol II molecules through nucleosomes. A) Metagene analysis of the MNase-seq data shows increased occupancy of the nucleosomes flanking the TSS and decreased occupancy of nucleosomes in the gene body in a *xrn1Δ* strain (in blue) compared to the wild type (orange). The red arrow indicates where the profile changes from higher nucleosome occupancy to lower occupancy in *xrn1Δ*. B) The same analysis as in A but around the polyA site (pAS) and dividing the genes depending on the gene length into 3 groups according to gene length. The red arrows indicate the intersection between wild-type and *xrn1Δ* profiles. C) Compiled BioGRO-seq signals for the wild type and *xrn1Δ* in +1, +2, +3 and +4 nucleosomes aligned to the dyad axis. Profiles of active RNA pol II were unchanged in the +1 nucleosome but were wider and shifted to the 3’ in *xrn1Δ* in the subsequent nucleosomes (+2, +3 and +4).

We also detected decreased occupancy of nucleosomes in the gene body, downstream of nucleosome +2 (Fig 2A). Metagenes obtained after aligning genes to the polyadenylation site (pAS) allowed the visualisation of nucleosome occupancy reduction in *xrn1Δ* along the whole gene body (Fig 2B). Independently of gene length, nucleosome occupancy of *xrn1Δ* started to be lower than the wild type around positions +400 (Fig S6) and remained depleted until the 3’ end of the gene body (Fig 2B). This is fully consistent with the alteration in transcription elongation deduced from the lower slope observed in RNA pol II CRAC profiles after Xrn1 perturbation. For instance, after 1h of nuclear Xrn1 depletion, the profile of RNA pol II CRAC progressively separated from the untreated sample decreased during the first 400 bp downstream of the TSS, independently of the gene length, and then remained constant until the pAS (Fig S7). This parallelism between increased density of RNA pol II molecules and reduced nucleosome occupancy is consistent with alterations of RNA pol II elongation dynamics caused by Xrn1 perturbations.

We wondered if the detected alteration in elongation would be reflected in the distribution of actively elongating RNA pol II molecules across the nucleosomes located in different positions of the gene body. We compiled the whole set of BioGROseq signals of +1 to +4 nucleosomes from the TSS, in order to get the average of each position in wild type and *xrn1Δ* cells, and represented them after alignment to the nucleosome dyad axis. We found no differences for the profile of +1 nucleosomes, suggesting that dynamics of the first nucleosome (usually occupying the TSS) during transcription initiation was not affected by lack of Xrn1 (Fig 2C). In contrast, we detected significant differences between *xrn1Δ* and the wild type for nucleosomes +2, +3 and +4. Higher density of BioGROseq signals were found before the dyad axis of these nucleosomes in the wild type, reflecting the accumulation of elongating RNA pol II just before the dyad of gene body nucleosomes. In *xrn1Δ* the profiles of actively transcribing RNA pol II molecules across +2, +3 and +4 nucleosomes were wider and shifted to the 3’, confirming a different dynamic of elongation immediately downstream of TSS (Fig2C).

### Xrn1 perturbation produces prominent pausing of RNA Pol II upstream of the pA site

RNA pol II CRAC also exhibited clear differences between wild-type and *xrn1Δ* cells at the 3’ end of genes. Prominent peaks were found at this region in *xrn1Δ*, usually located shortly upstream of the pA (Fig S8). Metagene analysis confirmed the presence of these 3’ peaks in different gene categories (Fig 3A, S9A-C), particularly in highly transcribed genes (Fig S9D-G). Albeit with different intensity, the vast majority of genes showed higher RNA pol II CRAC values in this region than the wild-type, with the maximal differential signal mapping 40 bp upstream of the pAS (Fig 3B-C).

**Figure 3.**
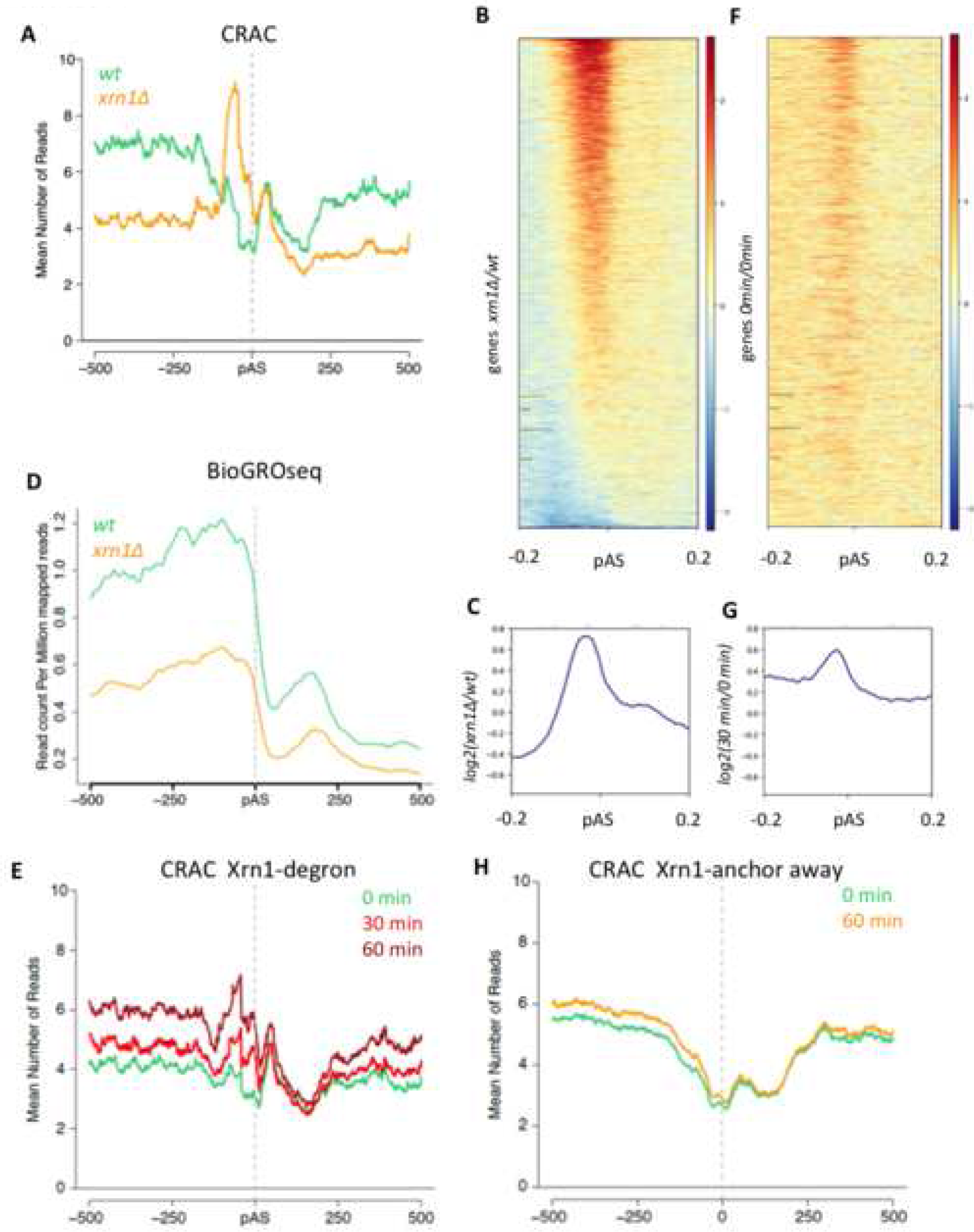
Xrn1 perturbation produces an alteration of total RNA pol II profiles at the 3’ end of genes. A) Metagene analysis of RNA pol II-CRAC profiles in *xrn1Δ* (orange) compared to the wild type (green) around the polyA site (pAS) of TATA-containing genes. B) Heatmap of the *xrn1Δ/*wt ratio of CRAC data, showing a widespread increase in the number of *xrn1Δ* reads just before the pAS (in red). Genes are ordered regarding the peak intensity. C) Metagene representing the mean of all genes shown in the previous heatmap. D) Metagene plot of the BioGRO-seq data in wild-type and *xrn1Δ* cells, showing that there is no accumulation of active RNA pol II in *xrn1Δ* at the 3’ end of TATA-containing genes. E) Metagene analysis of RNA pol II-CRAC profiles in Xrn1-Anchor away after 0 min (green), 30 min (red) and 60 min (burgundy) of Xrn1 depletion around the polyA site (pAS) of TATA-containing genes. F) Heat map of ratios, as in B, but 30 minutes after Xrn1 depletion (auxin-degron CRAC-seq data) divided by data before depletion. G) Metagene representing the mean of all genes shown in the previous heatmap. H) Metagene analysis of RNA pol II-CRAC profiles in Xrn1-Anchor away, showing no accumulation RNA pol II at the 3’ end of genes after nuclear Xrn1 depletion.

In contrast, no peak was found in this region in the BioGROseq profiles of *xrn1Δ* (Fig 3D). Since transcriptional run-on cannot detect backtracked RNA pol II molecules, we consider highly probable that the detected peak represents inactive molecules unable to move forward during very late elongation.

In order to confirm that the reads conforming the detected CRAC peaks corresponded to nascent RNA, all reads that contain a deletion (indicating the presence of a crosslink) were selected and a dataset containing only the position of the deletions (instead of the whole read) was generated. Genes showing prominent peaks were examined and showed patterns identical to those seen with all the reads (Figure S10). This accumulation of RNA pol II at the 3’ was also detected in metagenes after fast depletion of Xrn1 (Fig 3E, S11A), as it happened in almost every gene (Fig 3F, S10, S11B). We found, also in this case, that the maximal differential CRAC signal mapped 40 bp upstream of the pAS (Fig 3G, S11C). These results strongly indicate a functional influence of Xrn1 in late elongation, that is not due to the slow growth phenotype of *xrn1Δ*. However, we did not detect any RNA pol II CRAC signal accumulation at the 3’ end after depleting nuclear Xrn1 with the anchor-away technique, suggesting that this influence of Xrn1 on late elongation might not require its presence in the nucleus (Fig 3H).

Since the inactive RNA pol II peak at the 3’ does not seem to be directly related to nuclear Xrn1 but to lack of Xrn1 in the cell, we wondered if other perturbations of mRNA decay might produce similar results. We mapped total RNA pol II in *dhh1Δ* and *ccr4Δ*, two mutants lacking factors involved in decapping and deadenylation during mRNA decay, in a genome wide manner by ChIP seq. Metagenes showed increased levels of RNA pol II at 3’, around the pAS, in the two mutants, similarly to *xrn1Δ* (Fig S12A-C), suggesting that accumulation of RNA pol II at 3’ is a response to mRNA decay impairment. In all cases the intensity of RNA pol II accumulation increased with the transcription rate (Fig S12D-F).

We wondered whether the accumulation of inactive RNA pol II molecules at 3’ would involve alterations in the polyadenylation site selection. To test this, we processed raw data of poly(A) 3’T-fill experiments in *xrn1Δ* produced by Vicent Pelechano in Lars Steinmetz lab, who kindly provided them for this study. The overall distribution of mapped polyA sites in *xrn1Δ* was similar to the wild type, although a slight 20-30 bp shift to 3’ was detected (Fig S13A-B). However, the magnitude of this shift did not correlate to the intensity of RNA pol II accumulation at 3’, suggesting that these two phenomenon are not directly linked (Fig S13C).

### Late elongation of regulatory genes with poor nucleosome positioning at 3’ is particularly sensitive to lack of Xrn1

Transcription elongation defects can be due to impaired function of the elongation complex or to altered chromatin dynamics in front of elongating RNA pol II. Accumulation of inactive RNA pol II 40 bp upstream of the pAS implies that it takes place in the 3’ nucleosome-depleted region that characterizes RNA pol II-dependent genes. No significant alterations of nucleosome maps were detected in *xrn1Δ* cells in that specific position, in comparison to the wild type (Fig 2B). We also found no abnormalities in the main histone marks associated to the 5’ and 3’ chromatin dynamics during transcription. Histone H3 K4-trimethylation, a mark linked with initiation and early elongation, was not significantly changed in *xrn1Δ* in either a highly transcribed gene (*RPS3*) or a moderately expressed one (*SWD1*) (Fig S14A-B). Similarly, the profile of histone H3 K36-trimethylation, a modification linked to later steps of elongation, did not significantly change in any of the tested genes (Fig S14C-D).

Since we did not find any significant change of 3’ chromatin in *xrn1Δ*, we focused on the elongation complex. Transitions during transcription elongation are linked to modifications of the phosphorylation state of Rpb1 CTD, particularly in Ser2 residues (51). Tyr 1 phosphorylation has been also involved in termination (52). We analysed it by ChIP in three different genes. Two of them exhibited significant peaks in 3’ although with different transcription rates: *RPL40A* (highly transcribed) and *FLC2* (moderately transcribed). A third gene, *RPS3*, was selected for being highly transcribed, although it showed much less accumulation of RNA pol II at 3’ in *xrn1Δ*. Ser2 phosphorylation is the CTD mark most clearly associated to the elongation step of transcription. We found some alterations in the Ser2 profiles of the three genes tested in *xrn1Δ*, since higher levels of phosphorylation were detected in the mutant in all positions but the TSS (Fig 4B). In contrast, lower levels of Tyr1-phosphorylation was found in *FLC2* and in some amplicons of the two other genes tested (Fig 4C).

**Figure 4.**
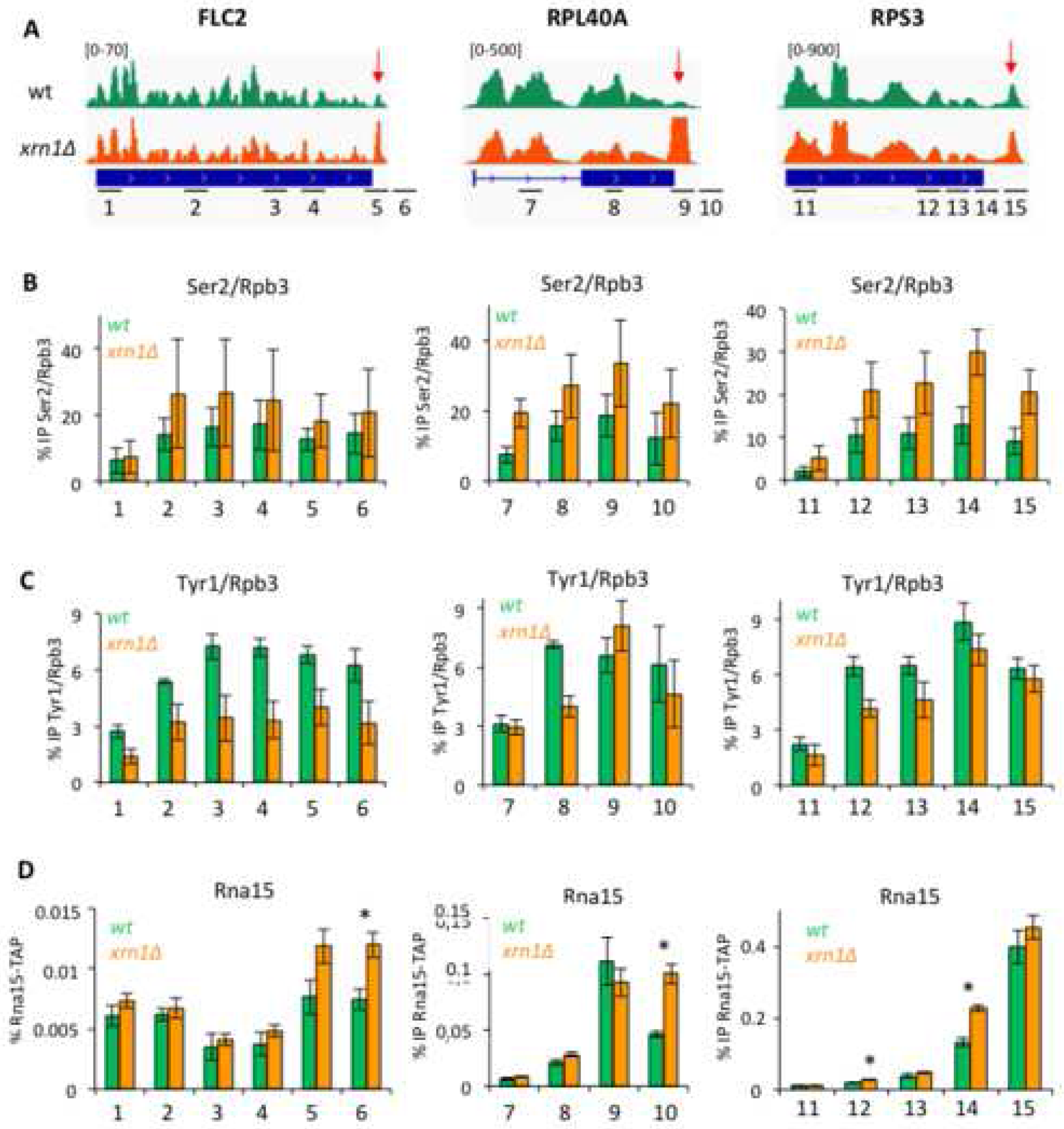
Lack of Xrn1 alters Ser2 and Tyr1 phosphorylation of RNA pol II CTD and enhances the binding of CF-IA. A) Snapshot of 3 different genes (from left to right: FLC2, RPL40A and RPS3) showing the CRAC signal in the wild type and *xrn1Δ* throughout the gene. The red arrow indicates the presence of an increased 3’ peak in the mutant. At the bottom we indicate with numbers the amplicons analysed in the subsequent ChIP experiments. B) Anti-phosphorylated Ser2-CTD ChIP experiment show slightly increased levels of this CTD modification in *xrn1Δ* strain in all three genes. We represent IP signal normalized by Rpb3 ChIP, obtained with the same extracts in parallel. C) Anti-phosphorylated Tyr1-CTD ChIP, normalized by Rpb3 ChIP, for the same genes. Slight decrease in Tyr1 phosphorylation was detected. D) Anti-Rna15-TAP ChIP shows increased binding of Rna15 to 3’ in *xrn1Δ* cells. In all cases the mean and standard deviation of three biological replicates is shown. All ChIP experiments were performed using the TAP-tagged Rna15 strain. * stands for a statistically significant difference between wt and mutant (Student’s t-test with p<0.05).

Ser2 and Tyr1 phosphorylation of Rpb1 CTD has been linked to the recruitment of 3’ pre-mRNA cleavage factors (51). Thus, we investigated whether lack of Xrn1 involved abnormal recruitment of CF-IA. We found significantly increased binding of Rna15 to amplicons located in the 3’ end of these and several other genes (Fig 4D and S15A-B). We also found increased levels of Rna14 at the 3’ end of *RPS3* (Fig S15C). These results indicate that in Xrn1-deficient cells the RNA pol II elongation complex is altered in two aspects related to 3’ mRNA processing: CTD phosphorylation and the binding of CF-IA. This suggests that the accumulation of inactive RNA pol II molecules at the 3’ end of genes might be the consequence of defective elongation complexes. However, this defect seems to be general and not specific of those genes showing the most prominent peaks.

Since the alterations in the elongation complex did not explain the intensity of 3’ peaks we wondered if the chromatin configuration of each gene might. We obtained metagene nucleosomal profiles of the 20% top and bottom genes in the distribution of peak intensities shown in Fig 3B. Clear differences were detected in the wild type (Fig 5A). Those genes with the most significant peaks showed a less defined nucleosomal pattern around the polyA site, and a wider and less-sharp nucleosome depleted region than bottom genes (Fig 5A). Similar differences between top and bottom genes were detected in *xrn1Δ*, indicating that these differences were neither enhanced nor suppressed by lack of Xrn1 (Fig S16).

**Figure 5.**
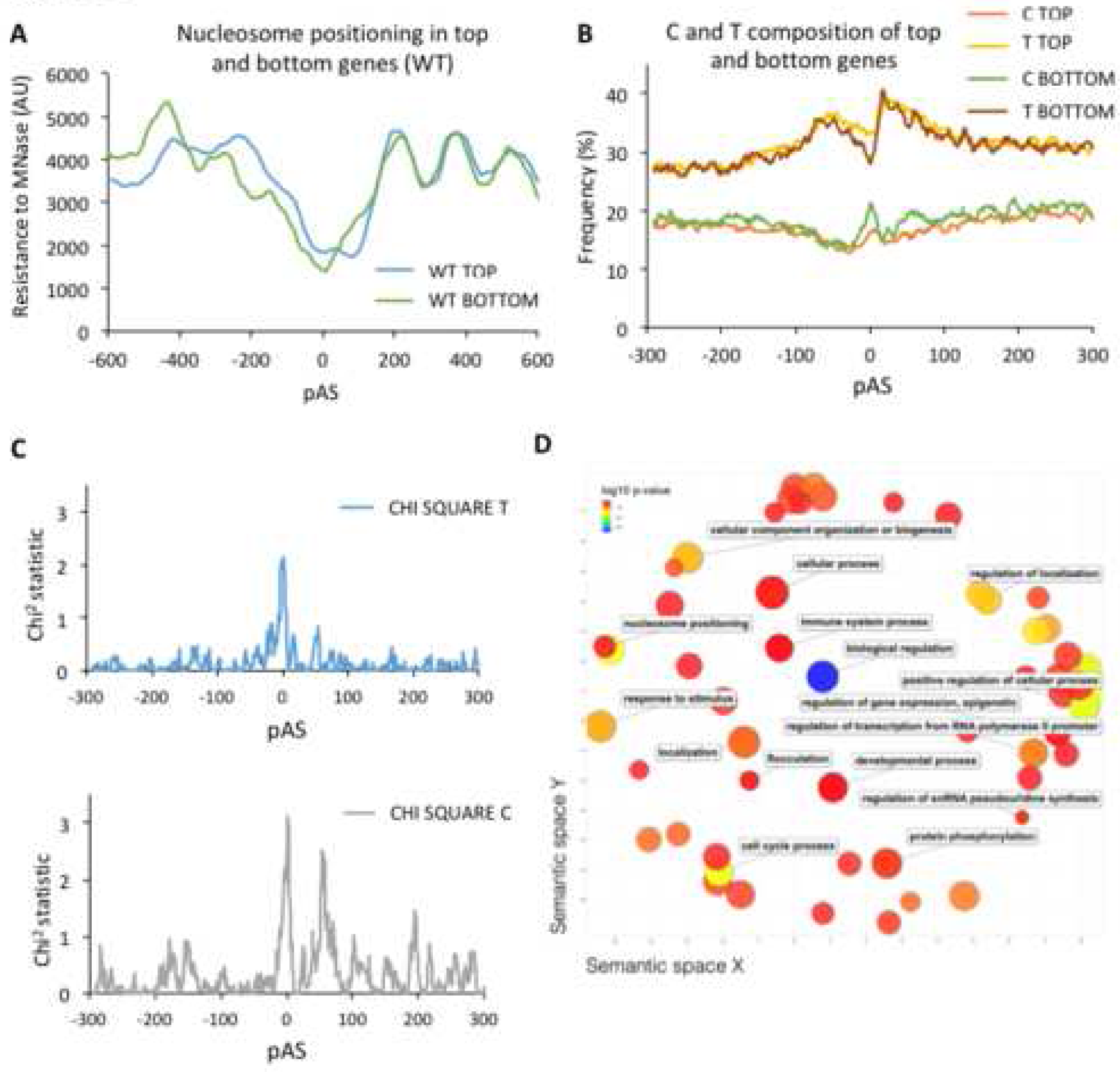
Genes with poor nucleosome positioning at 3’ in wild-type cells are more sensitive to RNA pol II arrest in *xrn1Δ*. A) Nucleosomal maps (in wild-type cells) of the pAS regions of genes showing different intensity of RNA pol II peaks at 3’ caused by *xrn1Δ*, obtained by resistance to micrococcal nuclease. Metagenes were calculated for two sets of genes from the distribution shown in Fig 3C: genes ranking in the 20% top (top genes) and 20% bottom (bottom genes). B) Frequency of C and T in the nucleotide composition of top and bottom genes. 10 nucleotide moving average is shown. C) Significance of the difference in C and T nucleotide composition between top and bottom genes, calculated by a Chi square test. D) Functional GO categories enriched in top genes.

We were unable to find any specific DNA motif differentially present in top genes versus bottom ones. However, the nucleotide composition of the pAS region exhibited some differences. Top genes showed a significant enrichment for T nucleotides in the pAS, whereas bottom genes exhibited higher frequency of C nucleotides in the same region (Fig 5C). This difference in sequence composition may explain the dissimilar nucleosomal organization of top and bottom genes around the pAS.

These results suggest that those genes with less positioned nucleosomes at their 3’ end are more sensitive to the alterations in late elongation caused by lack of Xrn1. In order to explore a possible functional consequence of this scenario we analysed the GO categories enriched in top genes. We found that most of the enriched GO terms were connected to the regulation of biological processes, suggesting that regulatory genes might be particularly sensitive to this phenomenon.

## Discussion

We have previously shown that lack of Xrn1 produced downregulation of transcription across the genome (1). In an independent work the group of E. Young also found a positive role for Xrn1 in the transcription of Snf1-dependent genes (53). However, a different study, based on RNA metabolic labelling, described apparently opposite effects of *xrn1Δ* on transcription (46). In this work, using different technical approaches, we demonstrate that Xrn1 perturbation originates significant negative alterations of RNA pol II distribution and, therefore, we confirm that Xrn1 plays a general positive contribution to gene transcription. Our experimental work suggests that metabolic labelling with 4-tU might introduce experimental noise in transcription rates calculations, particularly when the 5’ to 3’ pathway of mRNA degradation is impaired, as it happens in *xrn1Δ*. Previous results of Xrn1 were also based on the study of mutant cells lacking Xrn1, which exhibit pleiotropic phenotypes, including large cell volume and slow growth, which can indirectly affect transcription (48, 54). In this work, we made use of an auxin-degron approach to decrease the risks of indirect effects of *xrn1Δ*.

We also show in this study that Xrn1 perturbation specifically impairs transcription elongation at 5’ ends, shortly after initiation, and at 3’ ends, immediately before pre-mRNA cleavage. Similar observations have been very recently published after mapping RNA pol II distribution in *xrn1Δ* cells by NET-seq (55). At the 5’ end of gene bodies we observed that Xrn1 perturbations produced a change in the slope of the RNA pol II profiles obtained either with CRAC or BioGROseq. This change can be interpreted as the consequence of increased RNA pol II processivity (reduction in drop-off during elongation). This interpretation is contradictory with a positive role of Xrn1 in mRNA buffering, unless Xrn1 favored premature termination of non-productive elongating RNA pol II molecules. For instance, uncapped nascent RNA transcripts might be removed by Xrn1 during early elongation, like Rat1 does with nascent transcripts after pre-mRNA cleavage at 3’ (56).

Alternatively, the 50% reduction in slope produced by *xrn1Δ* can be interpreted as a decreased in RNA pol II elongation speed. This interpretation is in total agreement with the 50% reduction in RNA pol II elongation speed measured by us with the GAL1p-YLR454w system (31). It also fits with the enhanced recruitment of TFIIS across the genome observed in *xrn1Δ* cells, which indicates higher frequency of RNA pol II backtracking in the absence of Xrn1 (31).

We also found that nucleosome occupancy was decreased after the +2 position in the gene body, and that the high-resolution distribution of the BioGROseq signal across these nucleosomes was significantly altered in *xrn1Δ*. This parallelism between RNA pol II elongation defects and altered chromatin in gene bodies does not tell us whether there is a causative relation between them. However, these results shed light on the contribution of Xrn1 to chromatin transcription from two complementary aspects (nucleosome positioning and RNA pol II activity) and suggest a model were lack of Xrn1 makes the elongation complex unable to keep the wild-type elongation rate, due to different interaction dynamics between RNA pol II and the nucleosome. In this sense, Xrn1 might behave as other elongation factors that modulate RNA pol II pausing through nucleosomes (57).

At the 3’ end of gene bodies we observed the accumulation of RNA pol II CRAC signal 40 bp upstream of pAS in Xrn1-deficient cells. This indicates a strong pausing phenomenon. Lack of a similar signal in BioGROseq profiles, based on the run-on capacity of the mapped molecules, suggests that RNA pol II paused at this position become backtracked. This position of the gene body is a key point where the elongation complex undergoes severe changes. Some factors that travel with RNA pol II during elongation leave the complex at this point whereas others remain bound until termination (14). This restructuration is coupled to the 3’ processing of pre-mRNA, which is cleaved by CF-IA from the CTD. Arrest of RNA pol II at this point is likely reflecting abnormal dynamics of the elongation complex in this key step. Recruitment of CF-IA to the CTD during elongation is regulated by Ser2 phosphorylation (51). We found altered levels of these two CTD phosphorylation marks and enhanced binding of CF-IA to the elongation complex in Xrn1-perturbed cells. This suggests that the alteration of late elongation that we have detected is due to changes in the elongation complex that, later on, provokes RNA pol II arrest during the elongation/termination transition.

Although frequent, the peaks of RNA pol II shown by *xrn1Δ* at 3’ were not equally prominent in all genes. This specificity seems not to be related to the alterations of CTD phosphorylation marks or to the increased binding of CF-IA, since these changes also take place in genes with low accumulation of RNA pol II at 3’ gene ends. Our nucleosome maps indicate that these peaks are favored in genes with poor nucleosome positioning at 3’. So, it seems that these peaks result from the combination of altered elongation complexes in a certain chromatin environment. Intriguingly, genes with significant RNA pol II accumulation at 3’ are enriched in regulatory functions. This opens an interesting scenario, where genes with regulatory functions may reduce noise by mRNA buffering due to a particularly sensitive response to the mRNA decay/transcription crosstalk.

In principle, the two elongation phenotypes that we have described might be caused by a single molecular event derived from Xrn1 perturbation. However, several pieces of information suggest that they are partially independent and that only the alteration of elongation speed is directly related to a nuclear function of Xrn1. We have previously shown in individual genes that this function of Xrn1 is independent of Ccr4, as it is the genome-wide enhancement of TFIIS recruitment produced by *xrn1*≜ (48). The results of the Xrn1 anchor-away experiments must be interpreted cautiously (see above) but depletion of the nuclear Xrn1 pool seemed to confirm that Xrn1 early elongation function is related to its presence in the nucleus, whereas the late elongation function is not. In fact, accumulation of RNA pol II at 3’ was also detected in other mRNA decay mutants, like *dhh1Δ* and *ccr4Δ*, suggesting that the late elongation defect caused by lack of Xrn1 was mediated by its impact on mRNA decay. Impairment of mRNA degradation may cause an alteration in the circulation of factors between the cytoplasm and the nucleus, provoking a change in the composition of the RNA pol II elongation complex and/or in the nuclear availability of factors that regulate late elongation (see graphical abstract). This phenomenon would be particularly relevant for genes with poorly positioned nucleosomes at 3’. According to this view, Xrn1 would influence transcription elongation at least in two ways: by directly influencing elongation speed and by a more general mechanism that connects mRNA decay and late elongation.

## Accession numbers

All seq data are stored in the GEO repository (accession numbers GSE153037, GSE153072 and GSE158548).

To review GEO accession go to https://www.ncbi.nlm.nih.gov/geo/query/acc.cgi?acc=GSE153037 token: irixekwyrpkhnkf to https://www.ncbi.nlm.nih.gov/geo/query/acc.cgi?acc=GSE153072 token: cpkroosstvobdql and to https://www.ncbi.nlm.nih.gov/geo/query/acc.cgi?acc=GSE158548

### Acknowledgements

We thank Vicent Pelechano and Lars Steinmetz for generously sharing unpublished data, and M.E. Pérez-Martínez and M. Barneo-Muñoz for the BioGROseq setup. We also thank M. Choder for helpful discussion.

## Graphical abstract

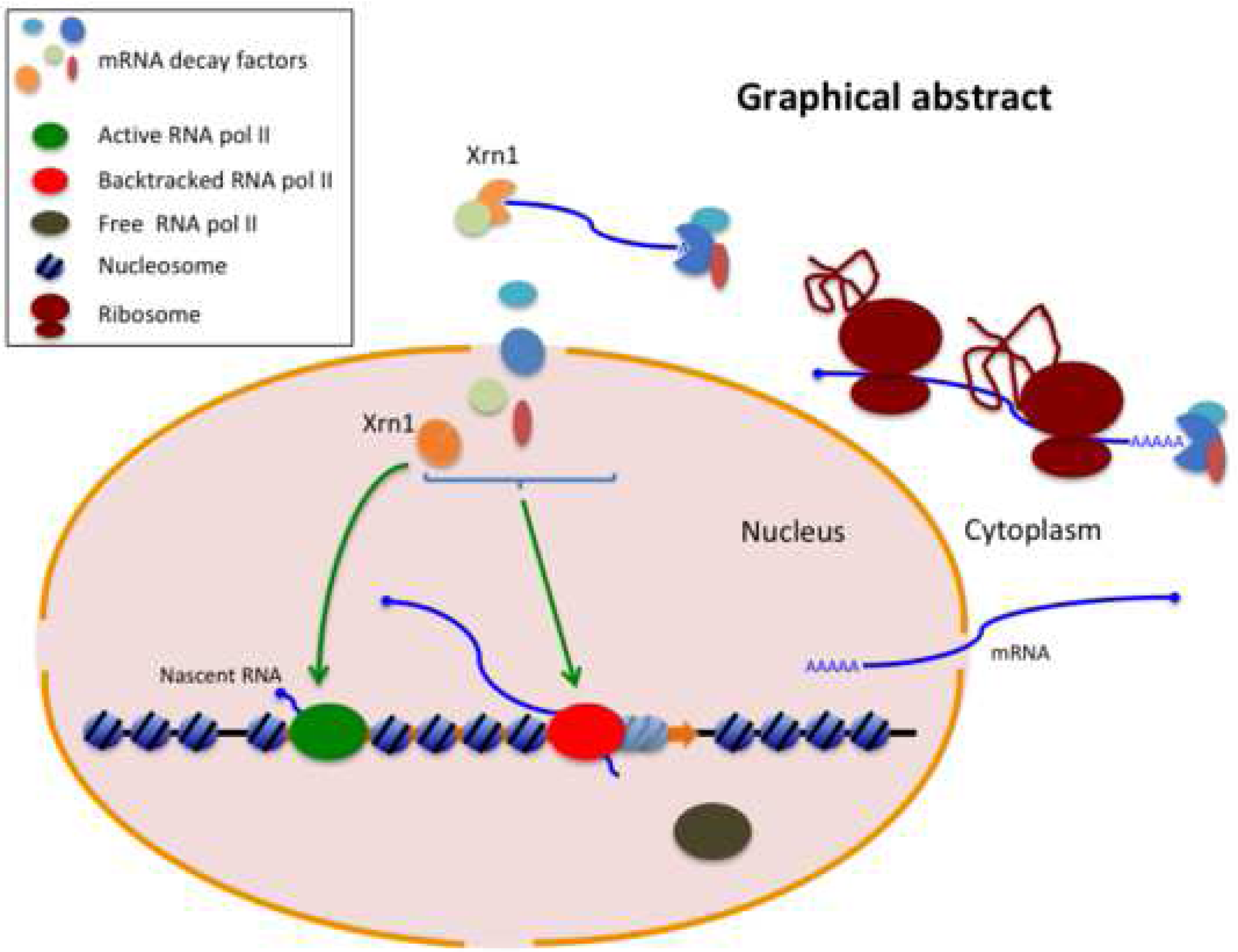

## Supplementary Figures

**Figure S1.**
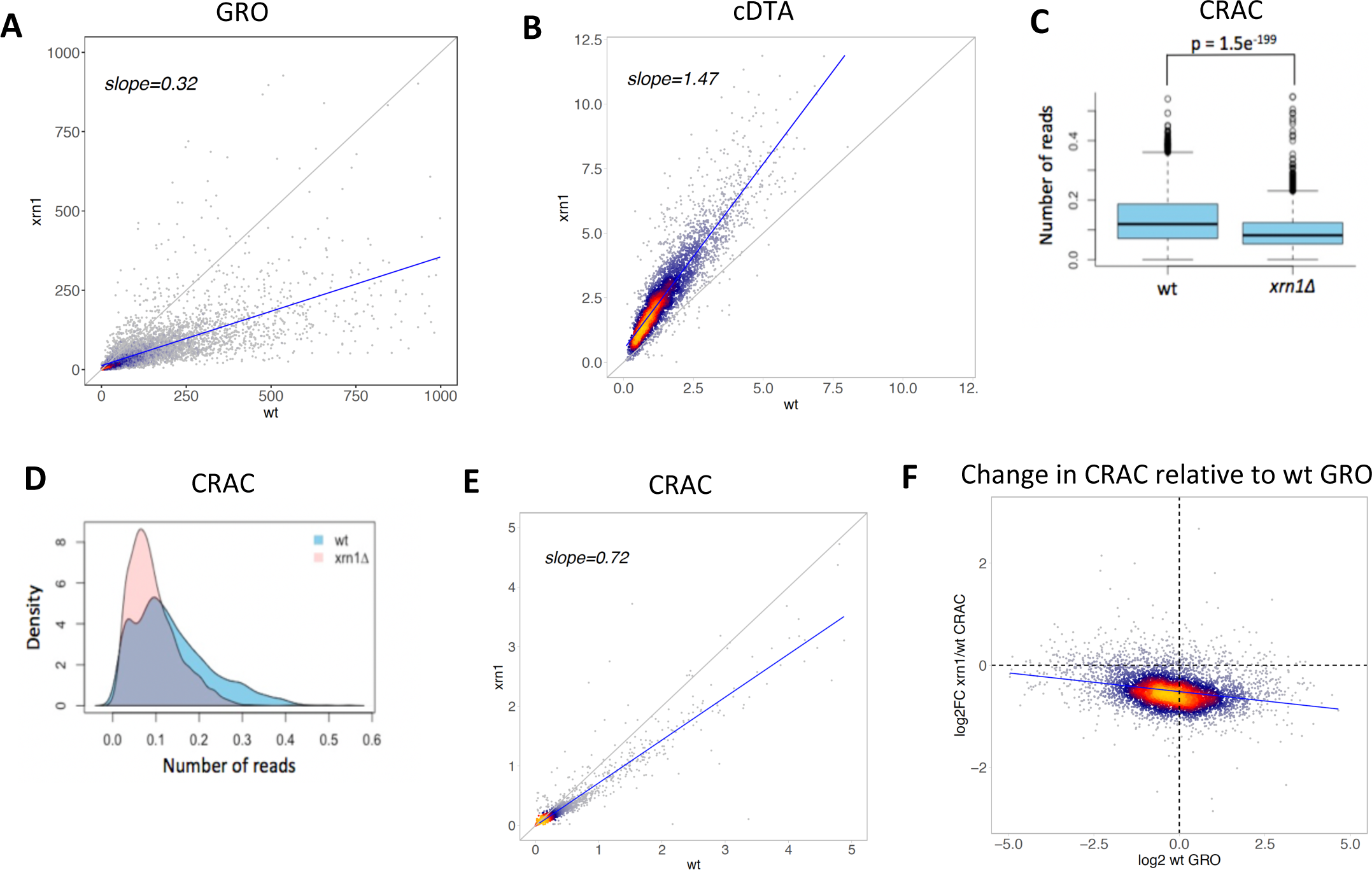
The deletion of *XRN1* led to a reduction in transcription using the CRAC and GRO methods. **A**. Scatterplot of the GRO data of *xrn1Δ* versus the wild type. We can see a clear down-regulation of most genes in the mutant as represented by the red regression line. The r=0.32 represents the slope of this regression line. The black line shows where the data should align if the two datasets had the same values. If the dots are under this line, then transcription is down-regulated in the mutant with respect to the wild type; and if they are above, then transcription is up-regulated. **B**. Scatterplot of the cDTA data obtained in Cramer’s lab (Sun et al., 2013). Most genes are up-regulated in the mutant compared to the wt as shown by the red regression line, with a slope of 1.47. **C**. Boxplot representing the number of reads normalized to gene length for the wild type and *xrn1Δ*. The black centre line represents the median of the number of reads (0.12 and 0.08 respectively). Then, reads are divided into quartiles. The box contains the reads from quartiles 2 and 3 (50% of all reads), and the whiskers show the quartile 1 (top whisker) and quartile 4 (bottom whisker). A two sample t-test shows that the differences are very significant (p<0.005) **D**. Density plot showing the density of reads versus the number of normalized reads. This represents how many times a read number appears in the data. **E**. Scatterplot of the CRAC data representing the number of normalised reads for the wild type versus *xrn1Δ*. Most genes are down-regulated in *xrn1Δ* in comparison to the wild type as represented by the red regression line with a slope of 0.72. **F**. Representation of the ratio between the CRAC signals of *xrn1Δ* and the wild type versus the wild type GRO data as a proxy for nascent transcription rate. The effect of *xrn1Δ* is greater in highly transcribed genes.

**Figure S2.**
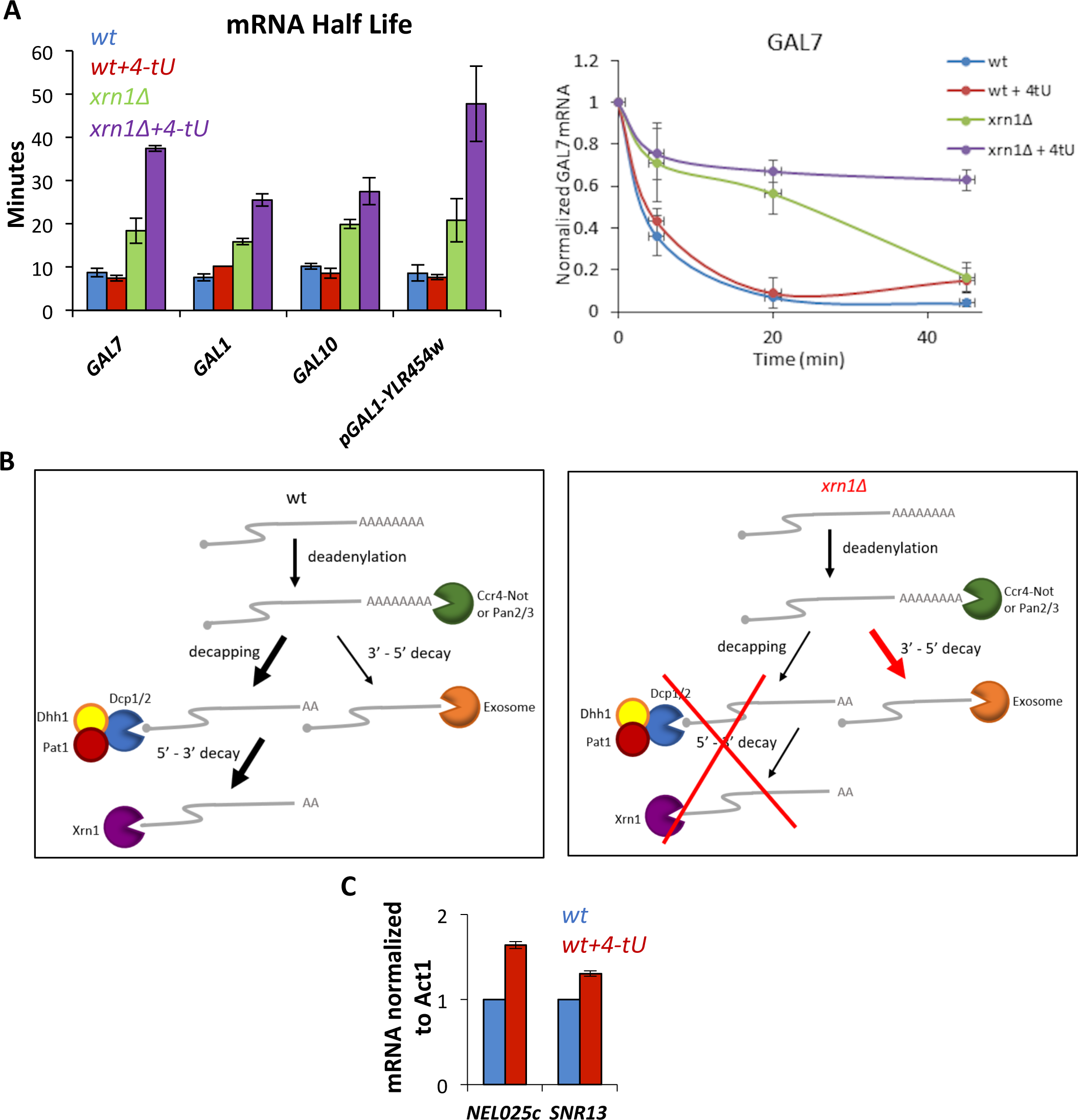
The use of 4-thiouracil (4-tU) leads to defects in degradation and transcription. **A**. Addition of 4-tU increases the mRNA half-life in an x*rn1Δ* strain. mRNA half-lives were estimated by using a transcriptional shut-off assay (see materials and methods), in a wt and x*rn1Δ* strain. Cells were grown to exponential phase and half of each type of cells were treated with 4mM 4-tU for 2 hours before performing the assay. mRNA signals were normalized to *SCR1*. The mean of two independent experiments are shown. Degradation kinetics of one of the mRNA analysed is shown on the right. **B**. Scheme showing mRNA degradation pathways in yeast (adapted from Braun and Young, 2014) in wt and x*rn1Δ*. **C**. Addition of 4-tU leads to an accumulation of ncRNAs in a wild type strain. Cells were grown to exponential phase and half of the cells were treated with 4mM 4-tU for 45 minutes. RNA signals were normalized to *ACT1*, an mRNA that is not degraded by the exosome. The mean of two independent experiments are represented by bars.

**Figure S3.**
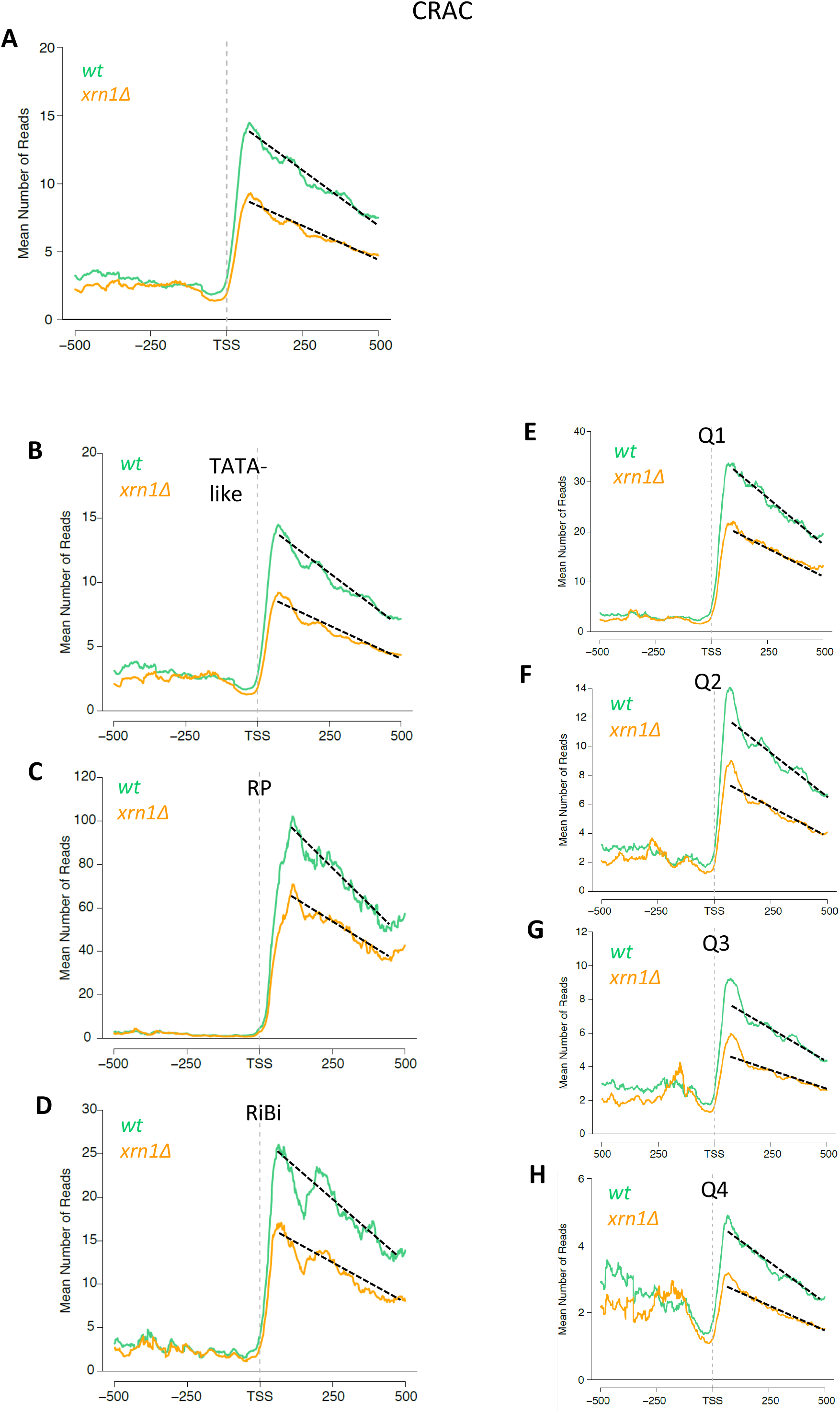
The transcription elongation defect in *xrn1Δ* is a widespread phenomenon. **A**. Metagene plot of the CRAC data of all genes aligned by their transcription start site (TSS). Compare the *xrn1Δ* strain (in orange) to the wild type (in green). **B-D**. In other gene categories we observe similar defects in *xrn1Δ* as for TATA genes (shown in fig 1A). We divided the genes into four categories: TATA (TATA promoter containing genes), TATA-like (TATA-like promoter genes, except for RP and RiBi genes), RP (ribosomal protein genes) and RiBi (ribosome biogenesis genes). **E-H**. In all quartiles there is a change in the slope in *xrn1Δ*.We divided the genes based on their transcription levels measured by CRAC into quartiles. Q1 is the quartile with the highest expressing genes and Q4 the lowest. This figure is related to Figure 1.

**Figure S4.**
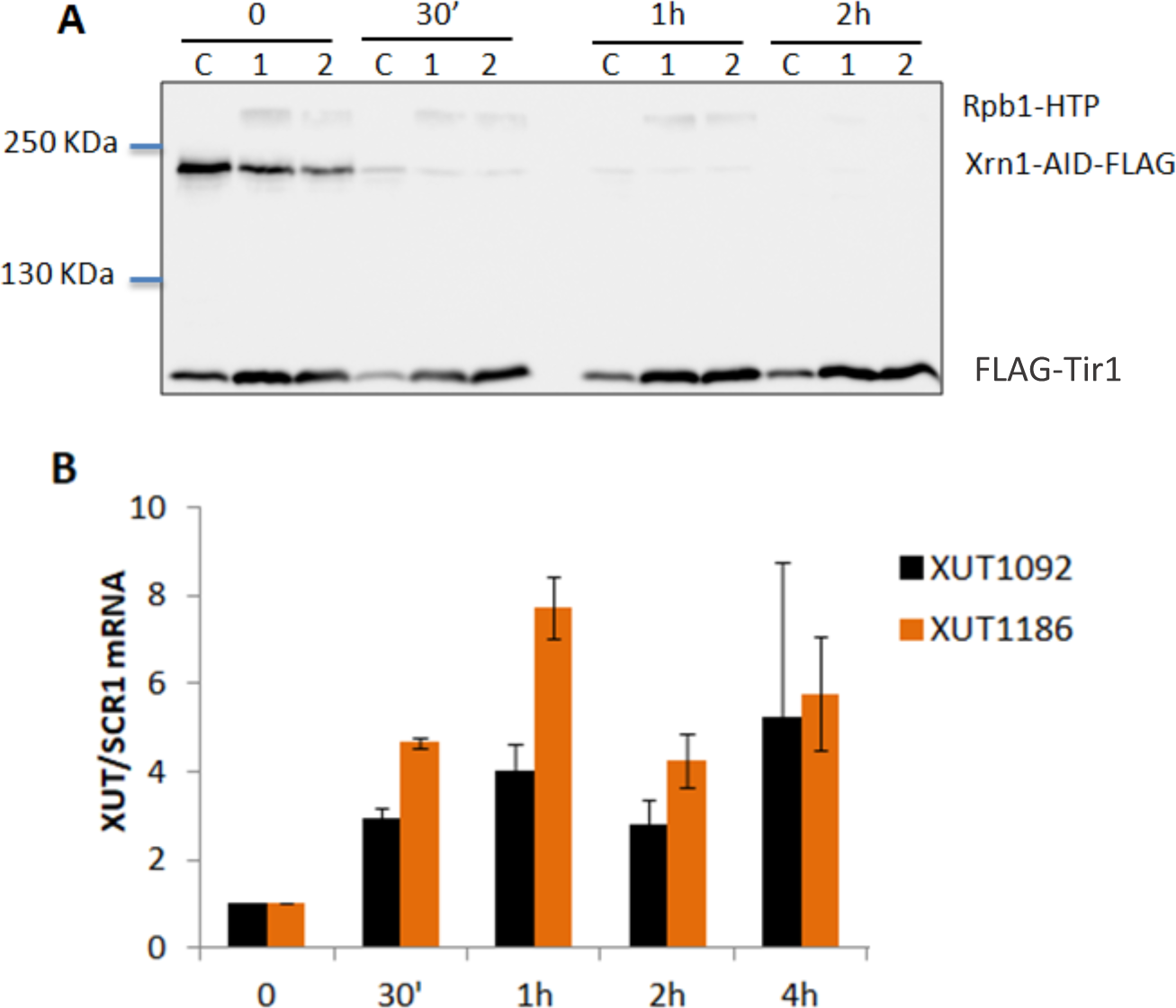
We successfully constructed a functional Xrn1-AID strain. **A**. Western blot showing the depletion of Xrn1-AID over time. The samples were obtained by growing cells to exponential phase and adding 0.2 mM of auxin and taking samples after the indicated times. The control strain, represented by a “C”, is the BY4741 strain used to transform our W303 Rpb1-HTP strain. The numbers 1 and 2 represent two tested clones obtained from the transformation. We incubated our membrane with an anti-flag antibody that allowed us to see Xrn1 and Tir1, which we used as control of protein amounts. The secondary IgG antibody used allowed us to also see Rpb1-HTP in the clones. **B**. RT-qPCR shows an increase in the expression of two XUTs (defined in Wery et al., 2016) after auxin mediated depletion of Xrn1. From here on out we used clone 1 in our experiments. Bars represent the average of two separate experiments. From these experiments we determined that a depletion of 30 minutes and 60 minutes were the best times to use for further experiments

**Figure S5.**
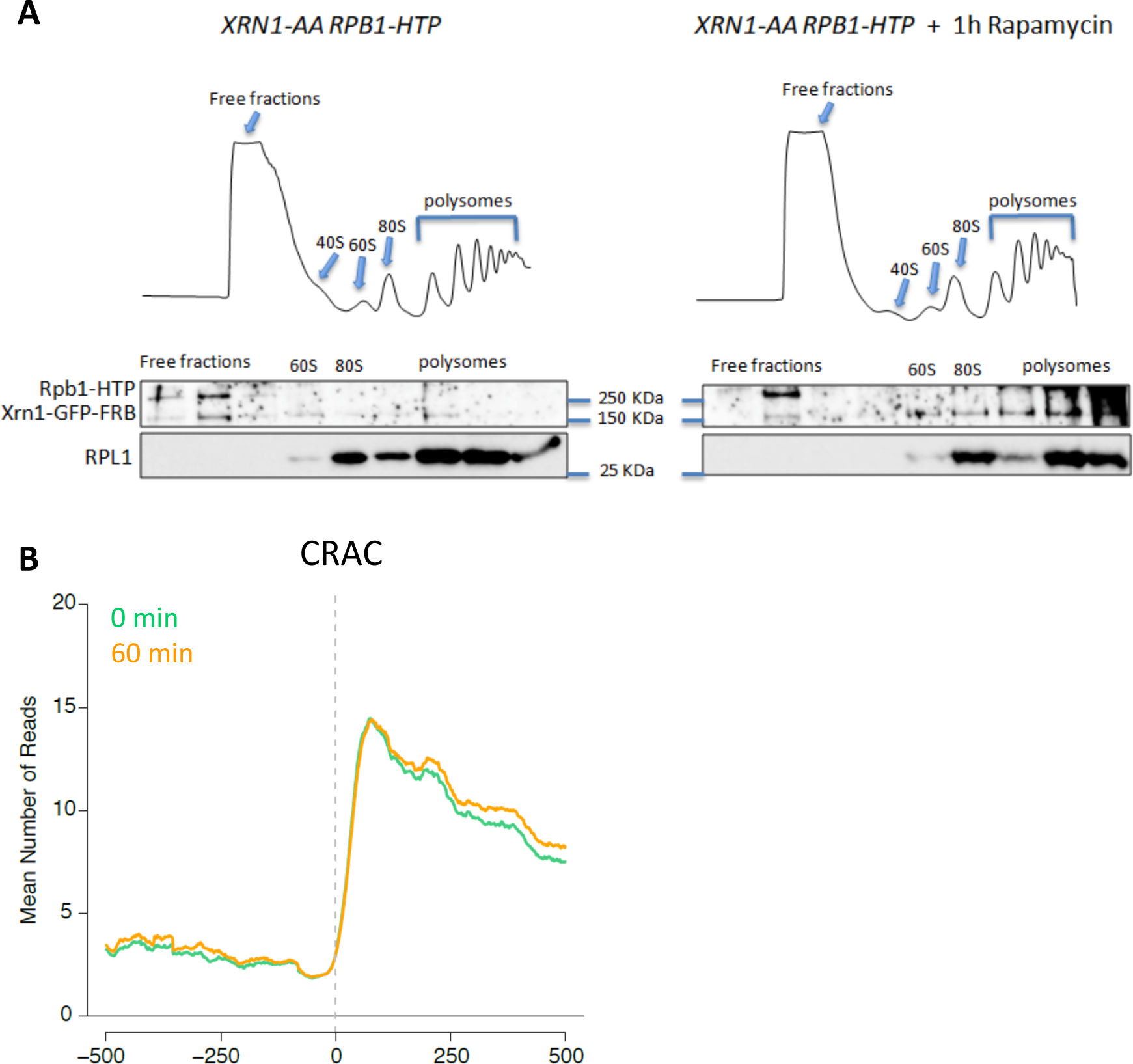
The Xrn1-AA strain is tethered to ribosomes upon rapamycin addition. We constructed the anchor away strain for our CRAC experiments from the strain used in Sun et al., 2013. **A**. Polysome profiles of the XRN1-AA RPB1-HTP strain with or without 1h rapamycin treatment. The fractions of these polysomes were collected and a western blot with the extracted protein was performed. We used anti-GFP to locate Xrn1, which also showed Rpb1-HTP, and anti-RPL1 to locate the RPL1 protein that assembles into the 60S ribosome subunit, 80S and polysomes. We show that without rapamycin, Xrn1 was mainly found in the free-fraction, and when rapamycin was added, Xrn1 was tethered to 60S ribosome subunits and polysomes. However, a part of the Xrn1 protein could still be found in the free fraction. **B**. Metagene analysis of Xrn1-AA aligning all genes to the TSS reveals early transcription elongation defects upon Xrn1 reduction in the nucleus. The green line shows the Xrn1-AA strain without rapamycin treatment (0 min) used as the control, and the orange line corresponds to 60 min rapamycin treatment.

**Figure S6.**
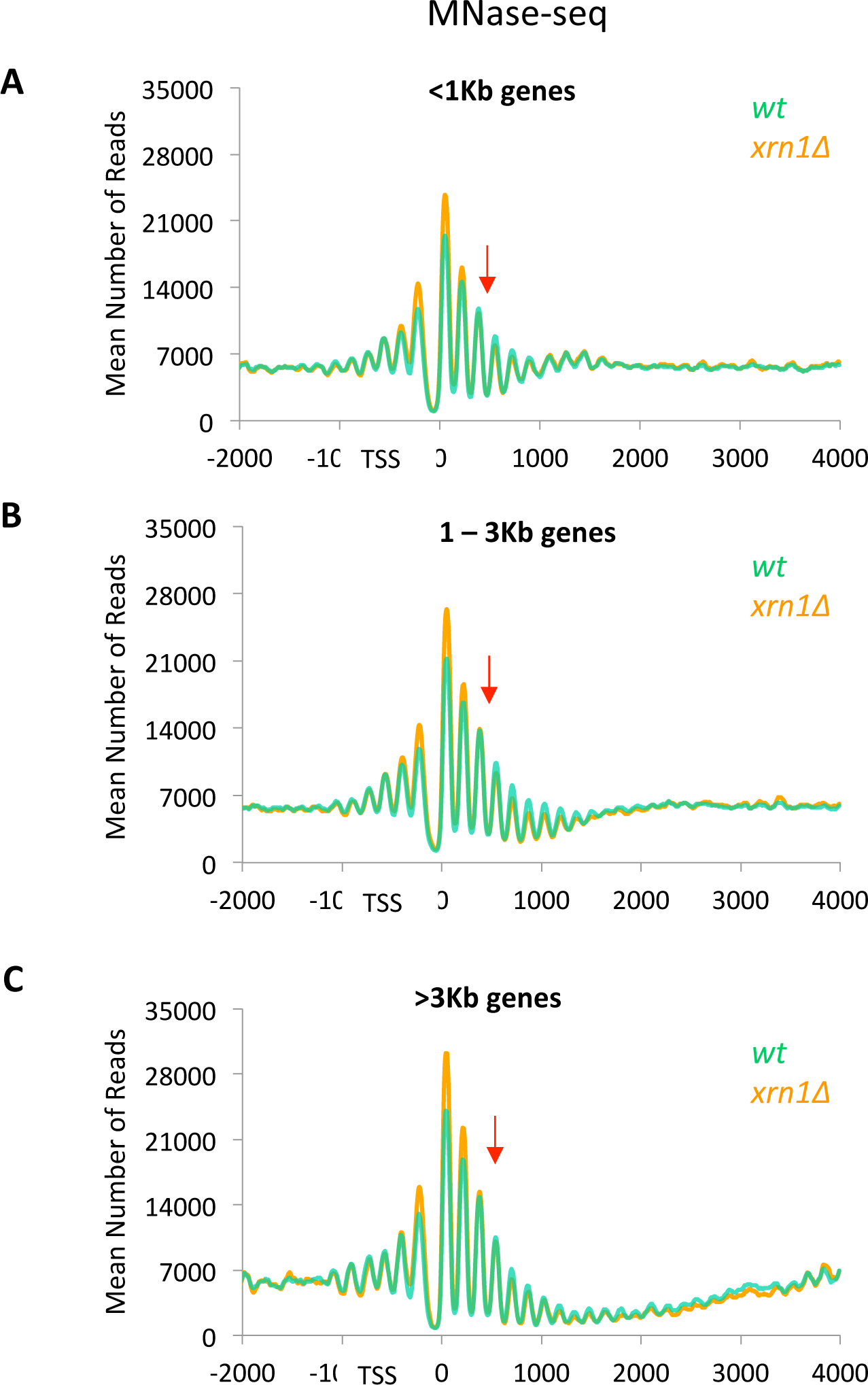
Changes in nucleosome occupancy was independent of gene length. We divided the genes by their length into three groups: less than 1 Kb (**A**), from 1 to 3 Kb (**B**) and longer than 3 Kb (**C**). We represented the nucleosomal maps of these genes aligned to the TSS. In green we show the wt and in orange *xrn1Δ*. As for all genes (Fig 2A), we detected in all cases a shift from higher nucleosome positioning to lower than the wt downstream of the +2 nucleosome. The red arrow indicates where the change occurs.

**Figure S7.**
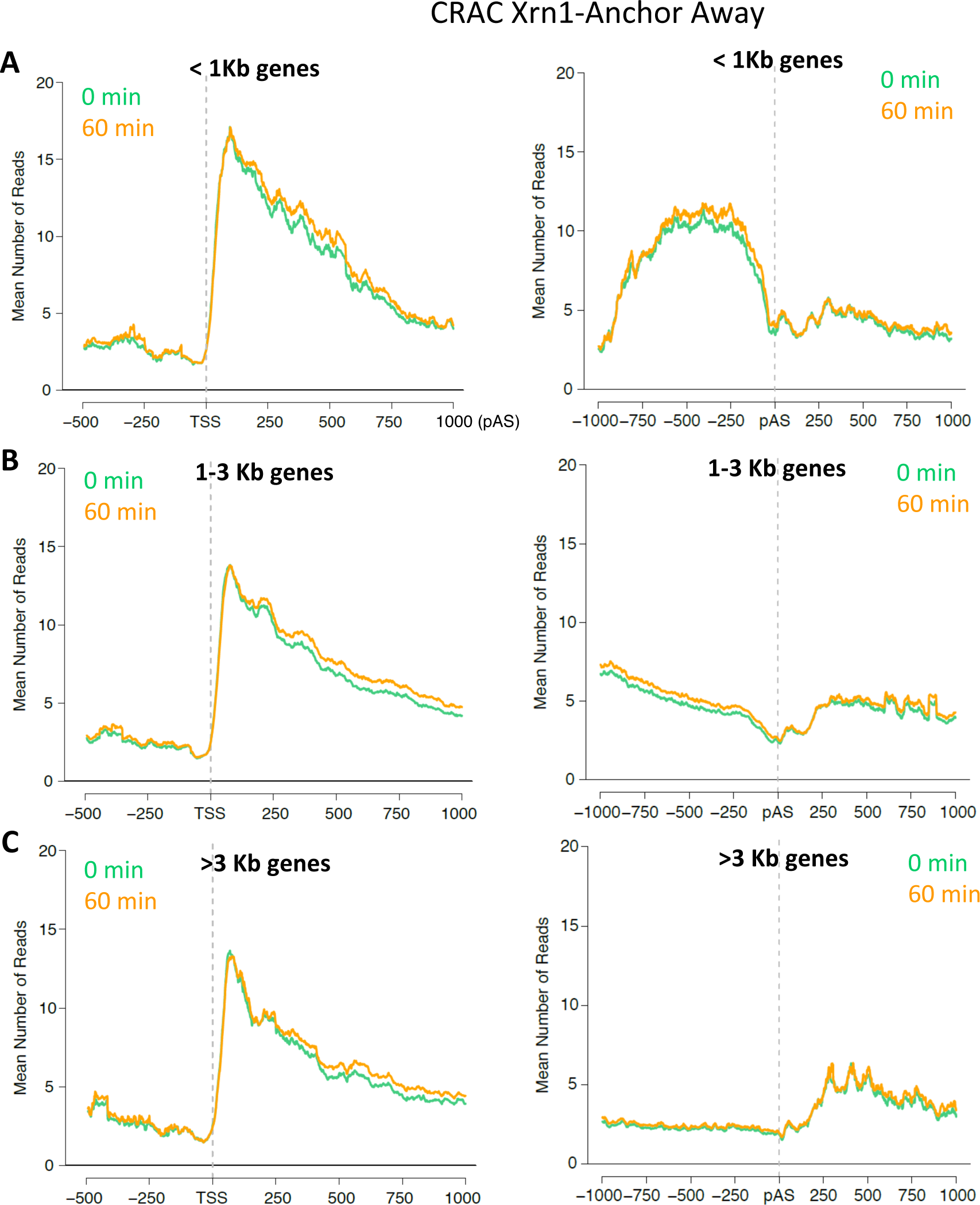
Metagene analysis of RNA pol II-CRAC data of Xrn1 AA experiment revealed as a function of gene length. The data used for Figure S6B were split into three groups according the length of their gene bodies (see Fig. 2B). **A**. Genes shorter than 1 kb. **B**. Genes between 1 and 3 kp long. **C**. Genes longer than 3 kb. No important differences between control and treated cells are seen for any gene length. In all cases the slope corresponding to the treated sample decreased with respect to the untreated sample in the 0-400 bp interval, and then the two continued in parallel.

**Figure S8.**
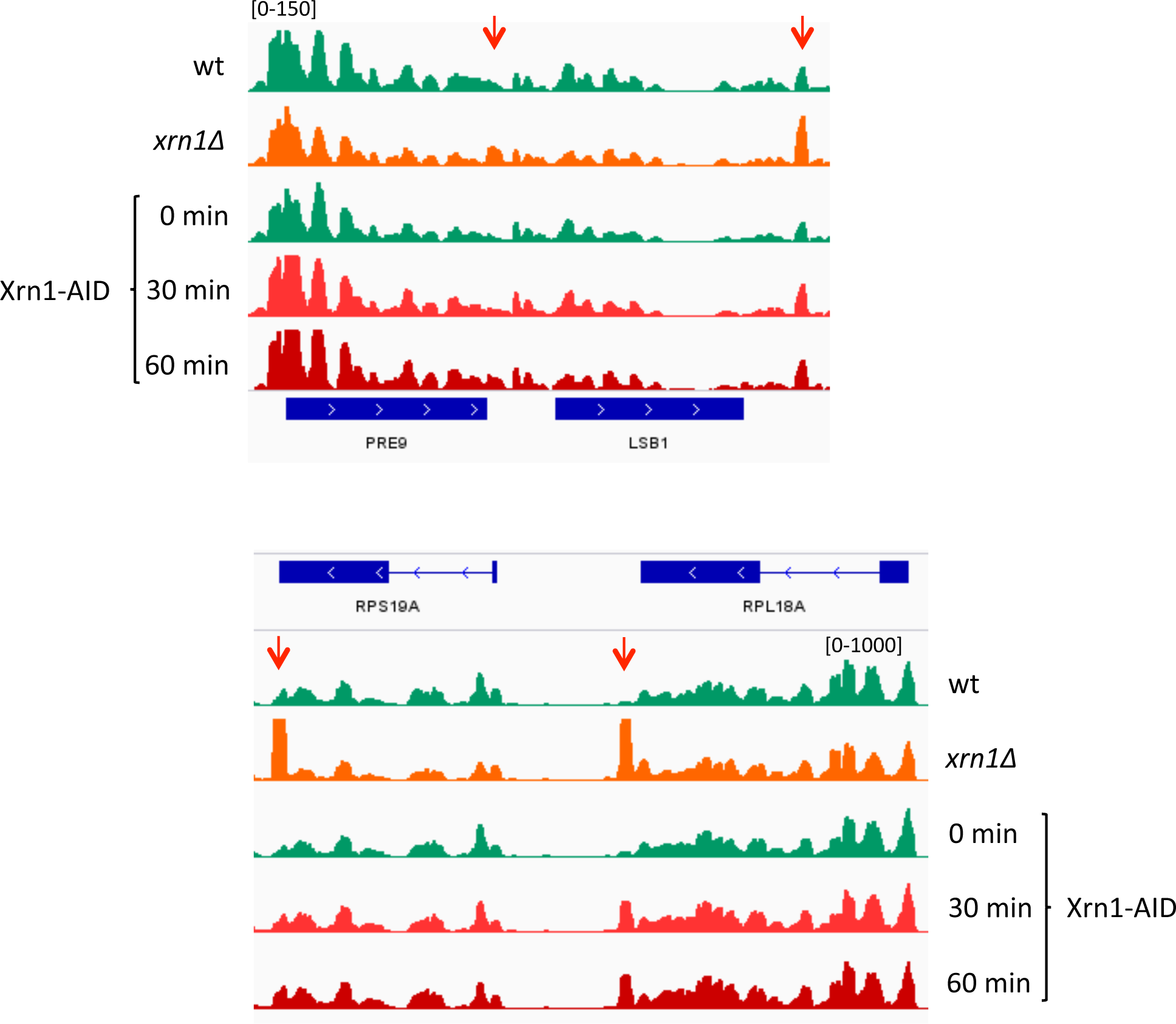
Several examples of genes showing RNA pol II peaks at 3’ in *xrn1Δ* and after depletion of Xrn1.

**Figure S9.**
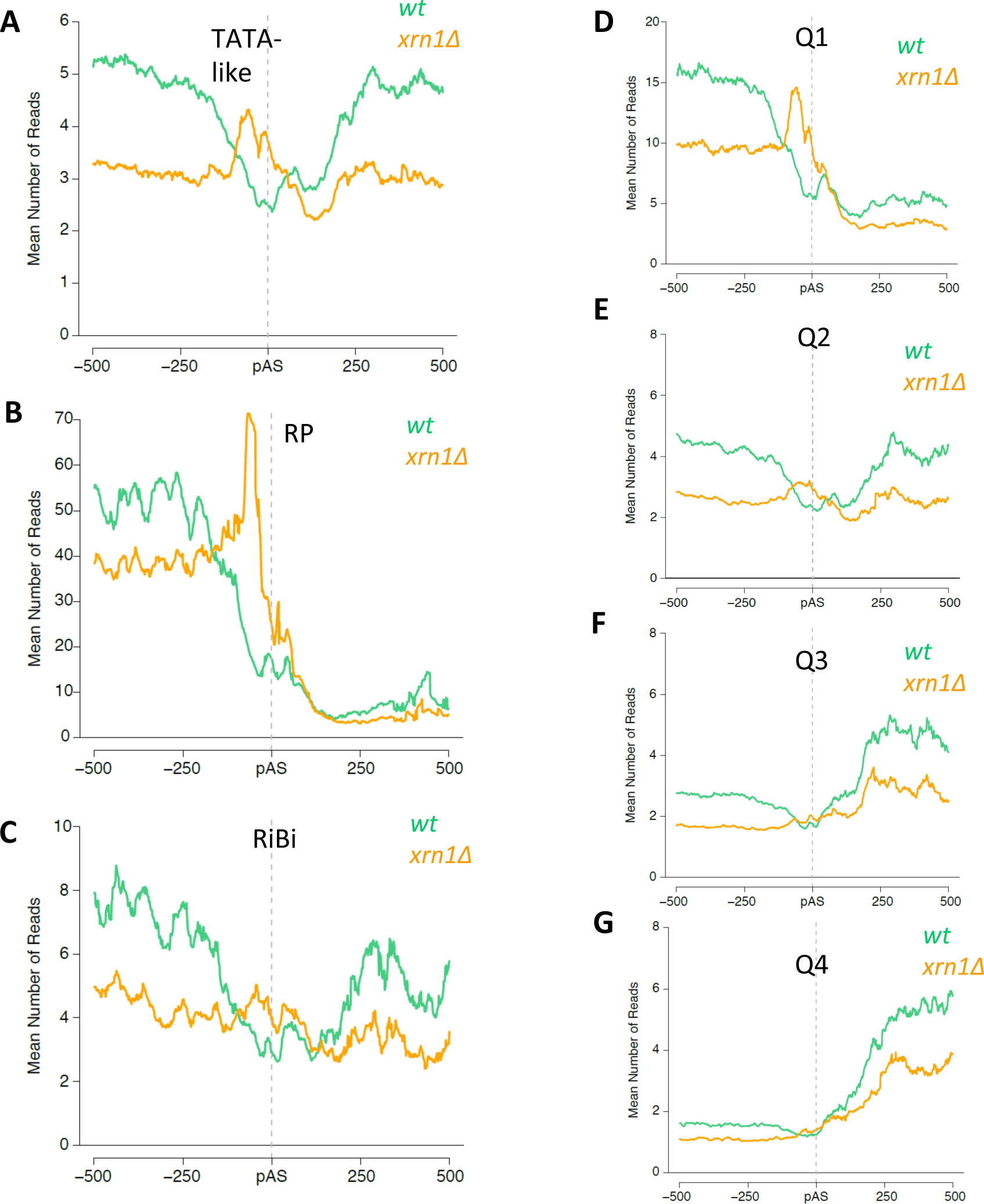
The late transcription elongation defect in *xrn1Δ* is very prominent in RP genes and specific of highly expressed genes. **A-C**. The accumulation of RNA pol II at the end of genes is very prominent in RP genes and detectable in TATA-like and RiBi genes. Genes were divided into categories (see fig S3) and aligned to their pAS. The green lines represent the wt and the orange lines the *xrn1Δ*. **D-G**. The accumulation of RNA pol II at the end of genes is very prominent in Q1, visible in Q2, but disappears in Q3 and Q4. Genes were divided into quartiles as in fig S3.

**Figure S10.**
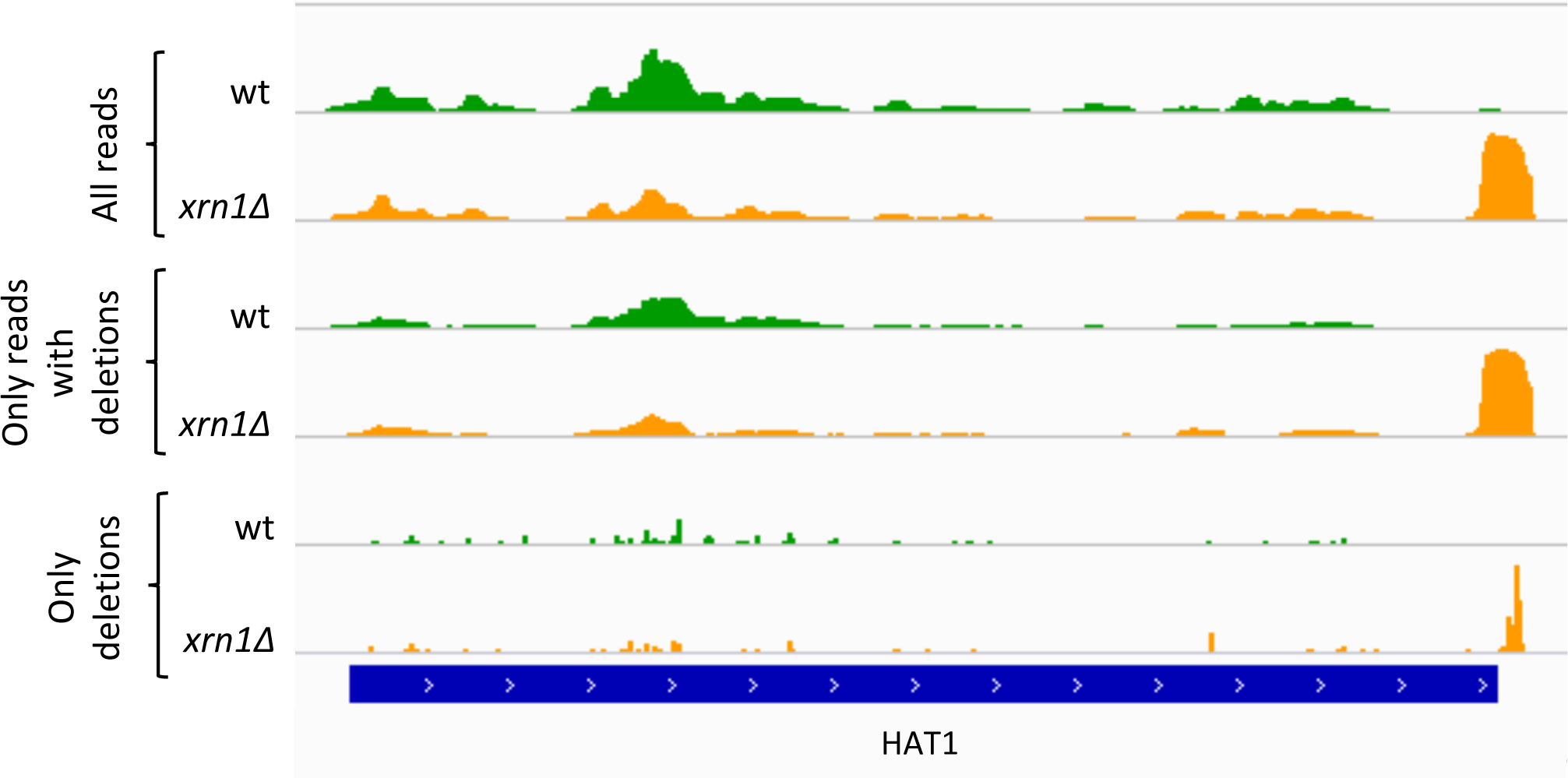
Deletion-containing reads confirm that 3’ peaks are conformed by elongating RNA pol II. All reads that contain a deletion (indicating the presence of a crosslink) were selected and a dataset containing only the position of the deletions (instead of the whole read) was generated. A gene showing a prominent peak of reads at 3’ and the corresponding peak of deletions is shown.

**Figure S11.**
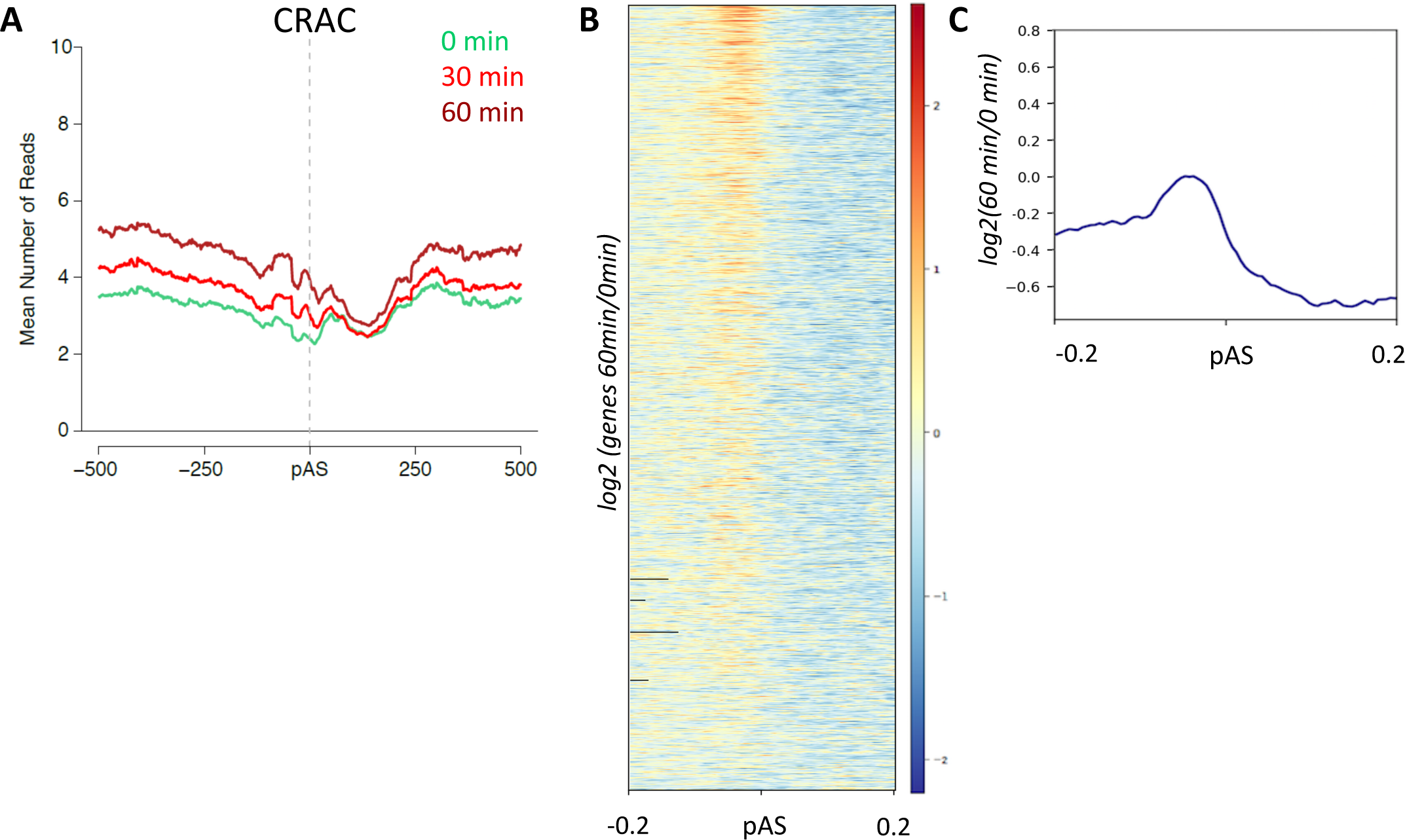
Depletion of Xrn1 using the degron system produced an alteration of total RNA pol II profiles at the end of TATA-containing genes. **A**. We found an increase in RNA bound to total RNA pol II measured by CRAC when depleting Xrn1 for 30 min (red) or 60 min (dark red) compared to the control (green) just before the pAS of genes. **B**. Heatmap of the Xrn1 depleted for 60 min data divided by the control data showing most genes present this increase in the amount of reads just before the pAS (in red). We show a +/- 200bp window from the pAS. Blue indicates lower reads, white no change, and in red higher amount of reads in the mutant versus the wt. **C**. Metagene representing the mean of what is shown in the previous heatmap.

**Figure S12.**
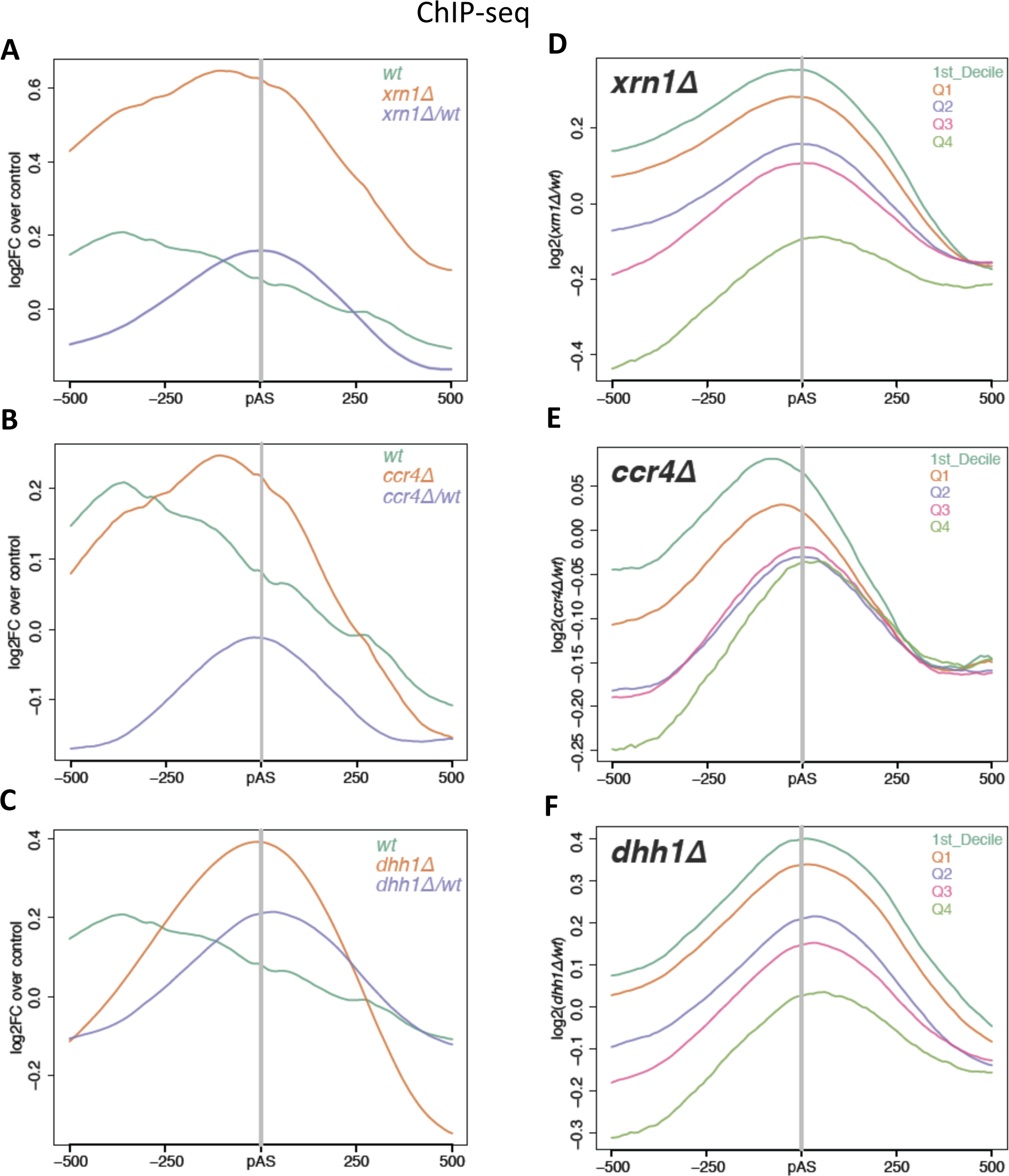
RNA pol II accumulates at the 3’ end of genes in mRNA decay mutants. **A**. Metagene analysis of Rpb3 ChIP-seq in *xrn1Δ* versus wt shows an increased accumulation of RNA pol II at the 3’ end of genes. In green we show the mean number of reads at each point of the wt (shown as IP/Input). In orange we see the result for *xrn1Δ*; and in purple we represent *xrn1Δ/*wt ratios. Metagene analysis of the Rpb3 ChIP-seq in *ccr4Δ* (**B**) and *dhh1Δ* (**C**) show similar results as for *xrn1Δ*. **D-F**. We show here the mutant/wt plots as in A-C but classifying the genes by their transcription rate. Decile 1 (D1) shows the top 10% of genes, quartile 1 (Q1) represents the genes with the highest signal and transcription rate, and quartile 4 (Q4) those with the lowest. The results show that the intensity of the pAS peak increases with the transcription rate.

**Figure S13.**
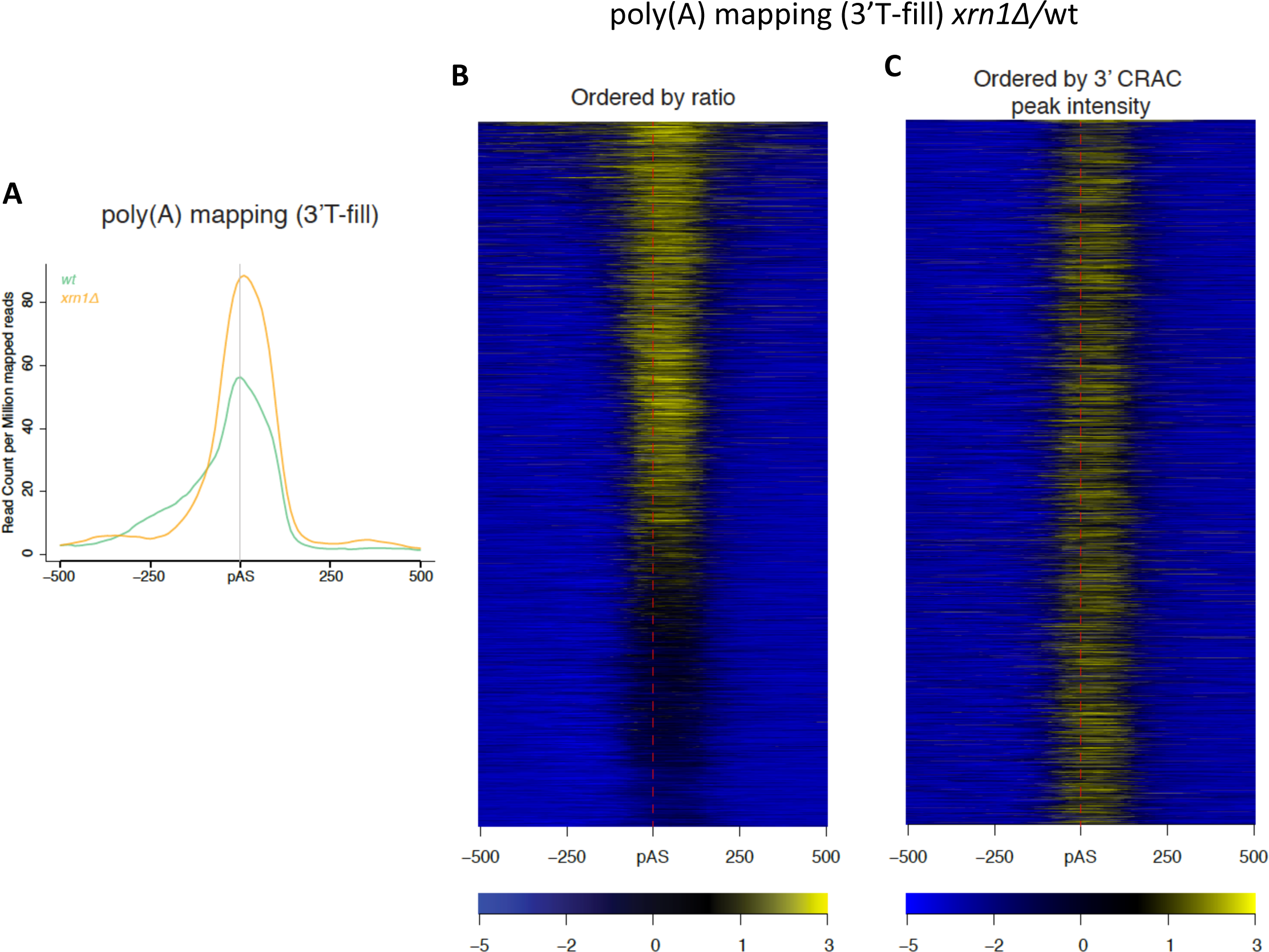
Lack of Xrn1 provokes minor downstream shifts of polyA sites but these changes do not correlate with the accumulation of RNApol II at 3’. **A**. Metagene showing the average normalised 3’T-fill read count around the poly(A) site (pAS) in wt and xrn1Δ. Te maximum density of 3’T-fill reads in wt aligns well with the main average annotated yeast pAS, whereas there is a downstream shift in xrn1Δ. **B**. Heat map of the log2FC xrn1Δ/wt 3’T-fill read density for individual genes around the main annotated pAS showing the downstream shift in pAS selection in xrn1Δ. **C**. Same heatmap with genes ordered according to the intensity of the 3’ peak of xrn1delta/wt ratio of CRAC data (related to Fig 3B).

**Figure S14.**
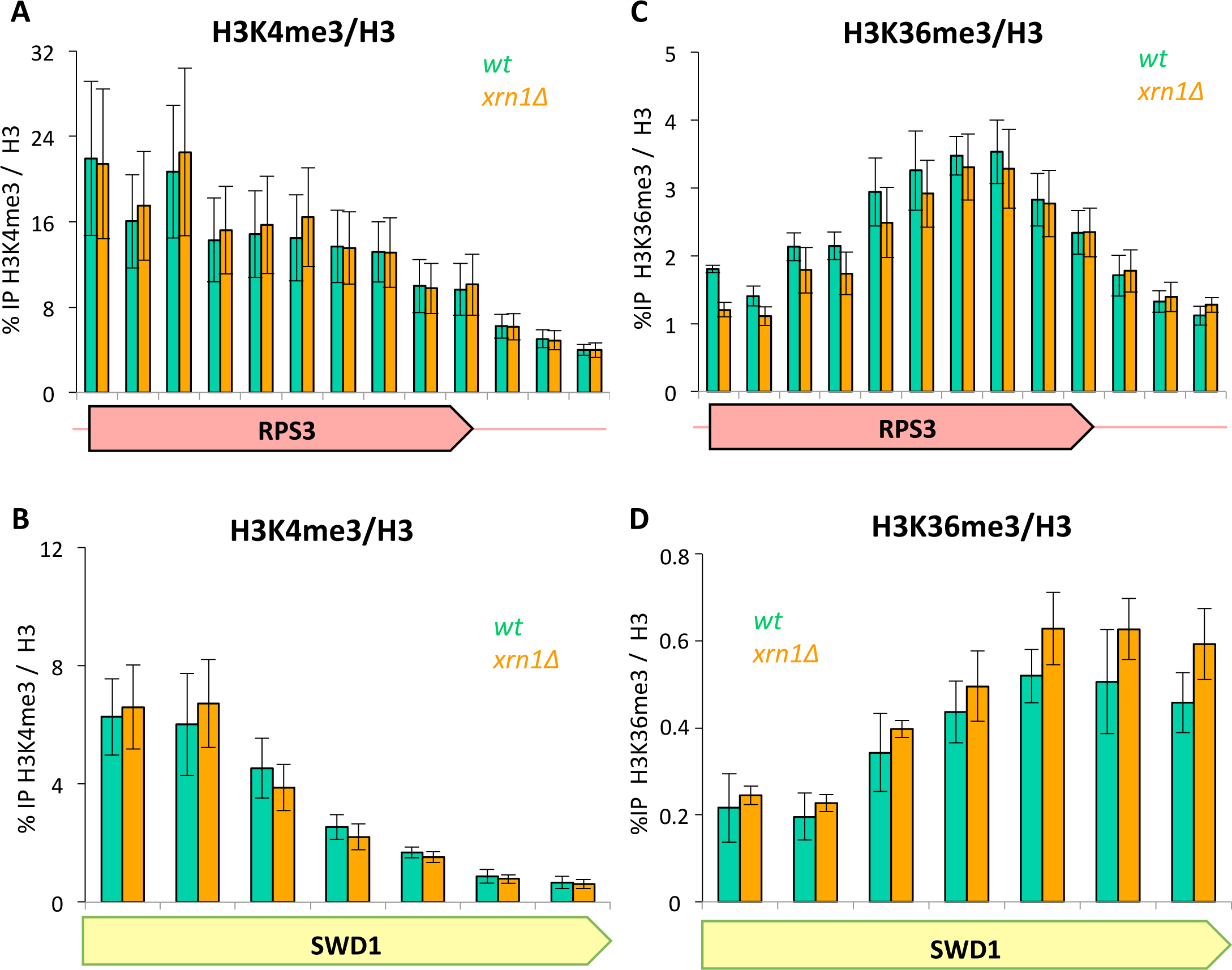
Changes in nucleosome patterns were not due to changes in histone modifications such as H3K4me3 and H3K36me3. **A**. No differences were found on the RPS3 gene between wt (green) and *xrn1Δ* (orange) in a H3K4me3 ChIP. the data was normalised to H3 to ensure changes were not due to changes in histone amounts, but in histone modifications (in this case methylation). **B**. No differences were found on the SWD1 gene between wt and *xrn1Δ* in a H3K4me3/H3 ChIP. **C**. No differences were found on the RPS3 gene between wt and *xrn1Δ* in a H3K36me3/H3 ChIP. **D**. No differences were found on the SWD1 gene between wt and *xrn1Δ* in a H3K36me3/H3 ChIP. In all cases the bars represent the mean values of three biological replicates.

**Figure S15.**
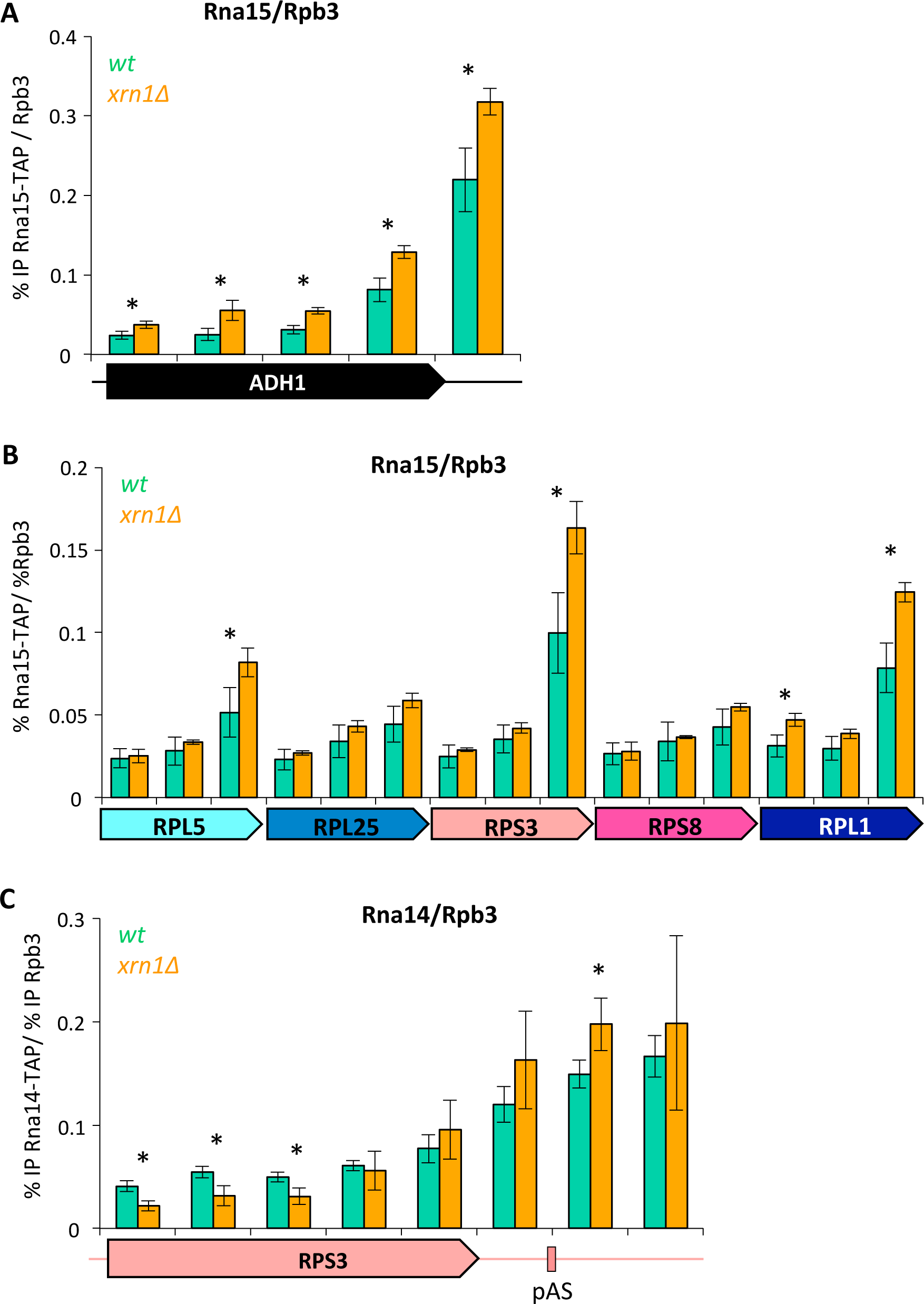
ChIP of Rna15/Rpb3 shows a significant increase of Rna15 on some genes in *xrn1Δ*. **A**. ChIP of Rna15/Rpb3 showed a significant difference on all analysed amplicons of the ADH1 gene. We performed the Rna15 ChIP in parallel to Rpb3 in order to normalize the Rna15 data and discard that any difference was due to differences in Rpb3 levels. We represent the mean signal (% IP) of three independent experiments along the ADH1 gene ORF (black arrow) and after (black line). The * shows a significant difference between wt and mutant using a Student’s t-test with p<0.05. **B**. We analysed three amplicons in five other RP genes and found a significant increase of Rna15 at the end of some of these genes (RPL5, RPS3 and RPL1). **C**. We also performed a ChIP of Rna14 (a protein tightly associated to Rna15) and also found some differences in *xrn1Δ*.

**Figure S16.**
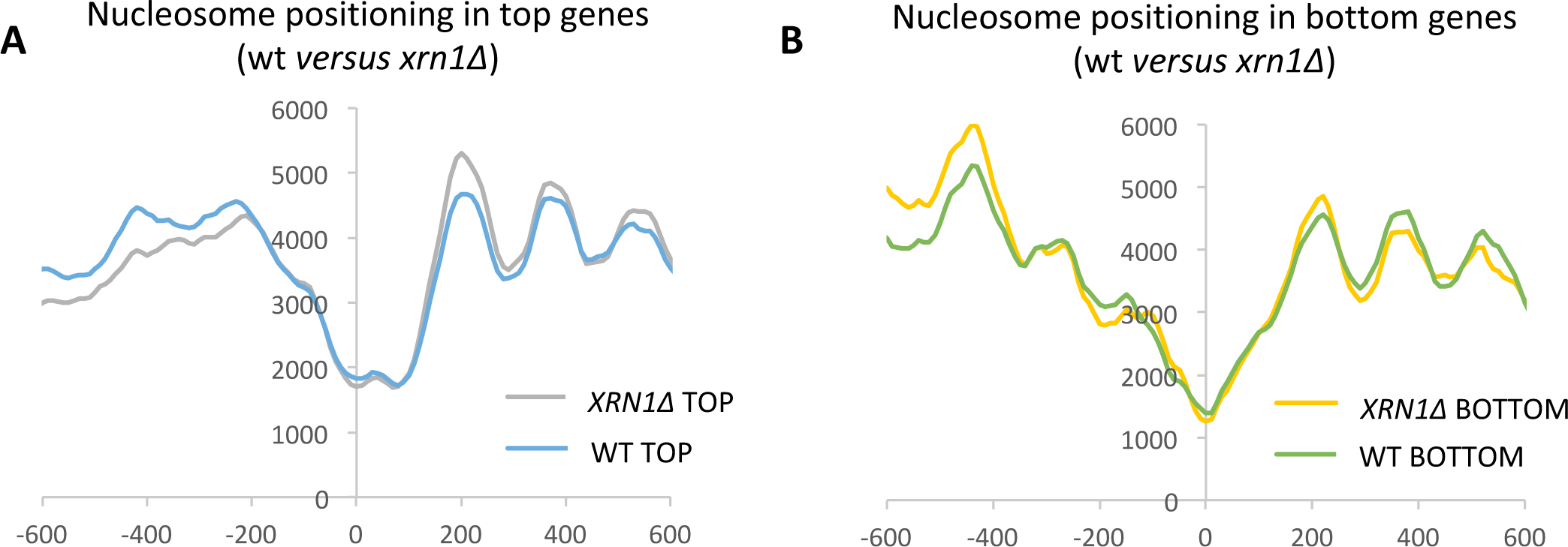
Nucleosomal patterns of top and bottom genes are not substantially altered by *xrn1Δ*. Maps of positioned nucleosomes of top (**A**) and bottom genes (**B**) around the pAS were obtained as an average of the patterns of resistance to MNase. Comparison of wt and *xrn1Δ* metagenes are shown.

## Supplementary Tables

**Table S1.** Yeast strains used in this study.

| *Yeast strain* | *Relevant genotype* | *Reference* |
| --- | --- | --- |
| RPB1-HTP | *W303 RPB1-HTP::TRP* | D. Libri Lab |
| VBY70 | *W303 RPB1-HTP::TRP Xrn1::KanMx4* | This work |
| VBY67 | *W303 XRN1-FRB-GFP::kanMx4 RPBL13-2xFKBP12::TRP tor1-1 frp1::LEU RPB1-HTP::URA* | This work |
| VBY72 | *W303 XRN1-AID-FLAG::KanMx4-TIR1 RPB1-HTP::TRP* | This work |
| Xrn1-AID | *BY4741 XRN1-AID-FLAG::KanMx4-TIR1* | D. Libri Lab |
| wt | *BY4741* | Euroscarf |
| *xrn1Δ* | *BY4741 Xrn1::KanMx4* | Euroscarf |
| *ccr4Δ* | *BY4741 Ccr4::KanMx4* | Euroscarf |
| VBY129 | *BY4741 Xrn1::KanMX4 Ccr4::KanMx4* | This work |
| VBY83 | *BY4741 Rna15-TAP::His* | Euroscarf |
| VBY90 | *BY4741 Rna15-TAP::His Xrn1::KanMx4* | This work |
| VBY82 | *BY4741 Rna14-TAP::His* | Euroscarf |
| VBY86 | *BY4741 Rna14-TAP::His Xrn1::KanMx4* | This work |

**Table S2.**
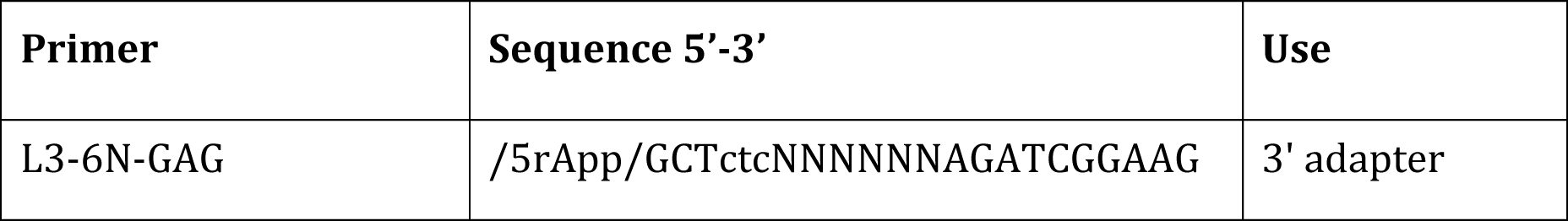
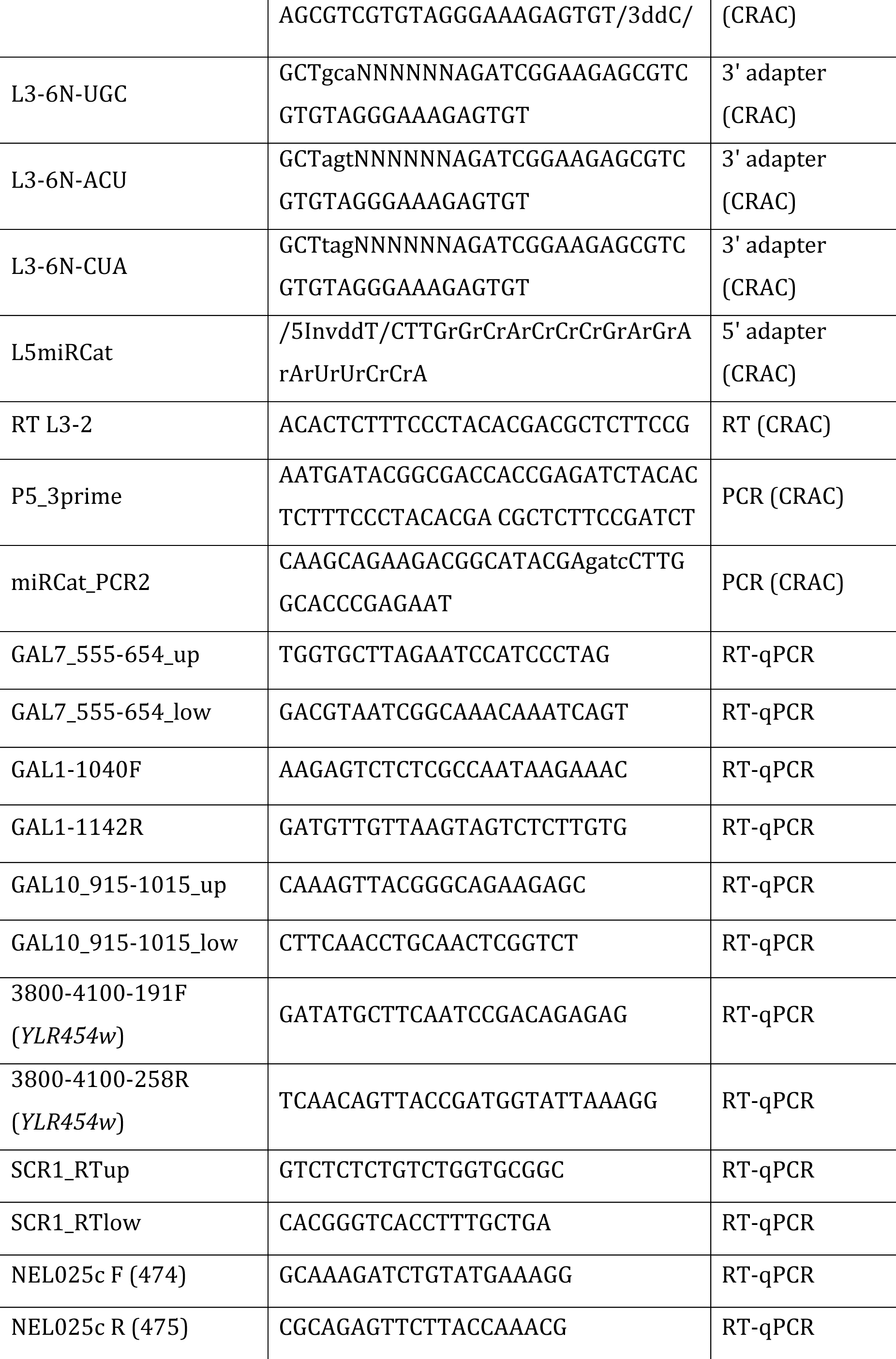
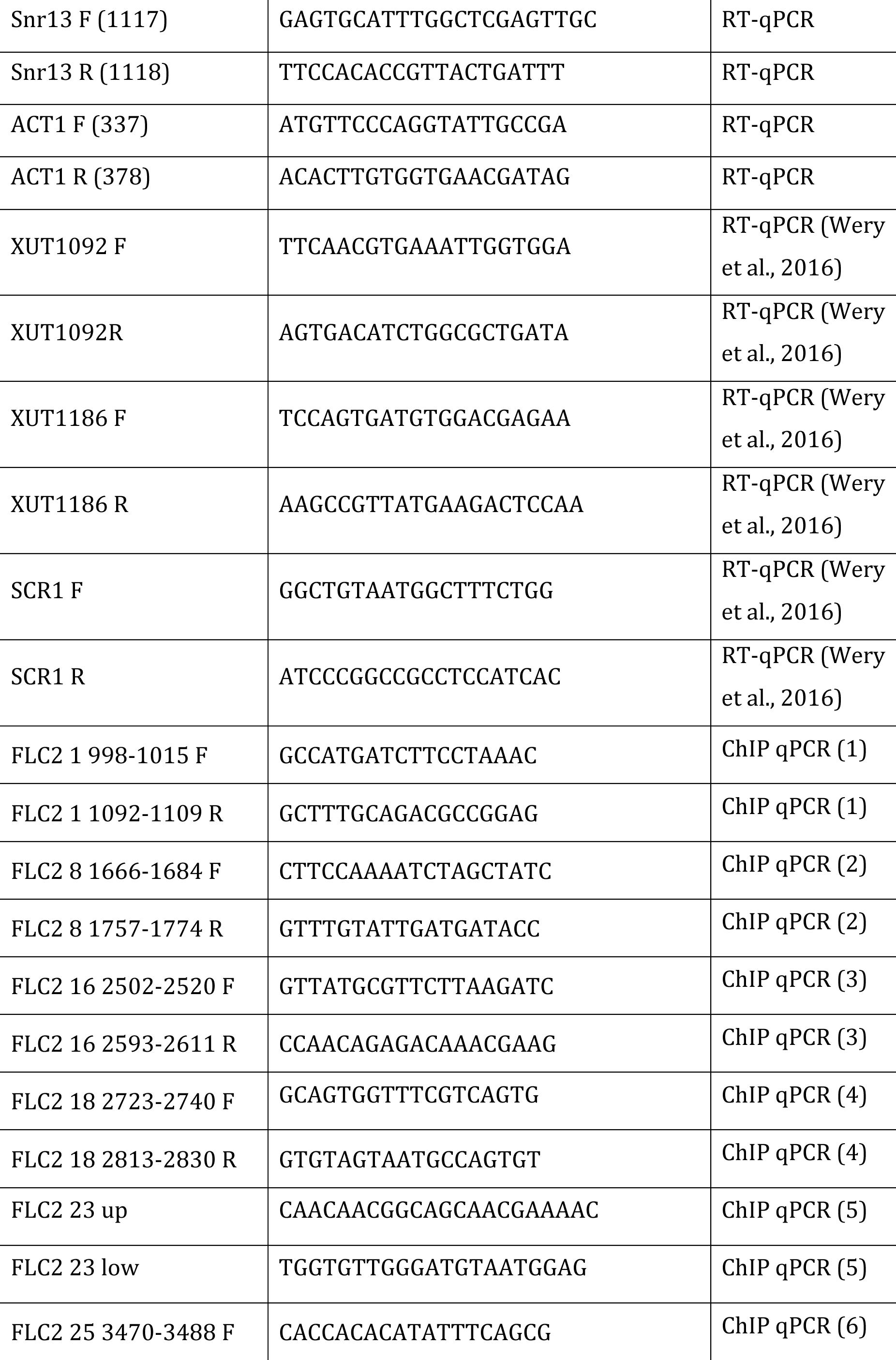
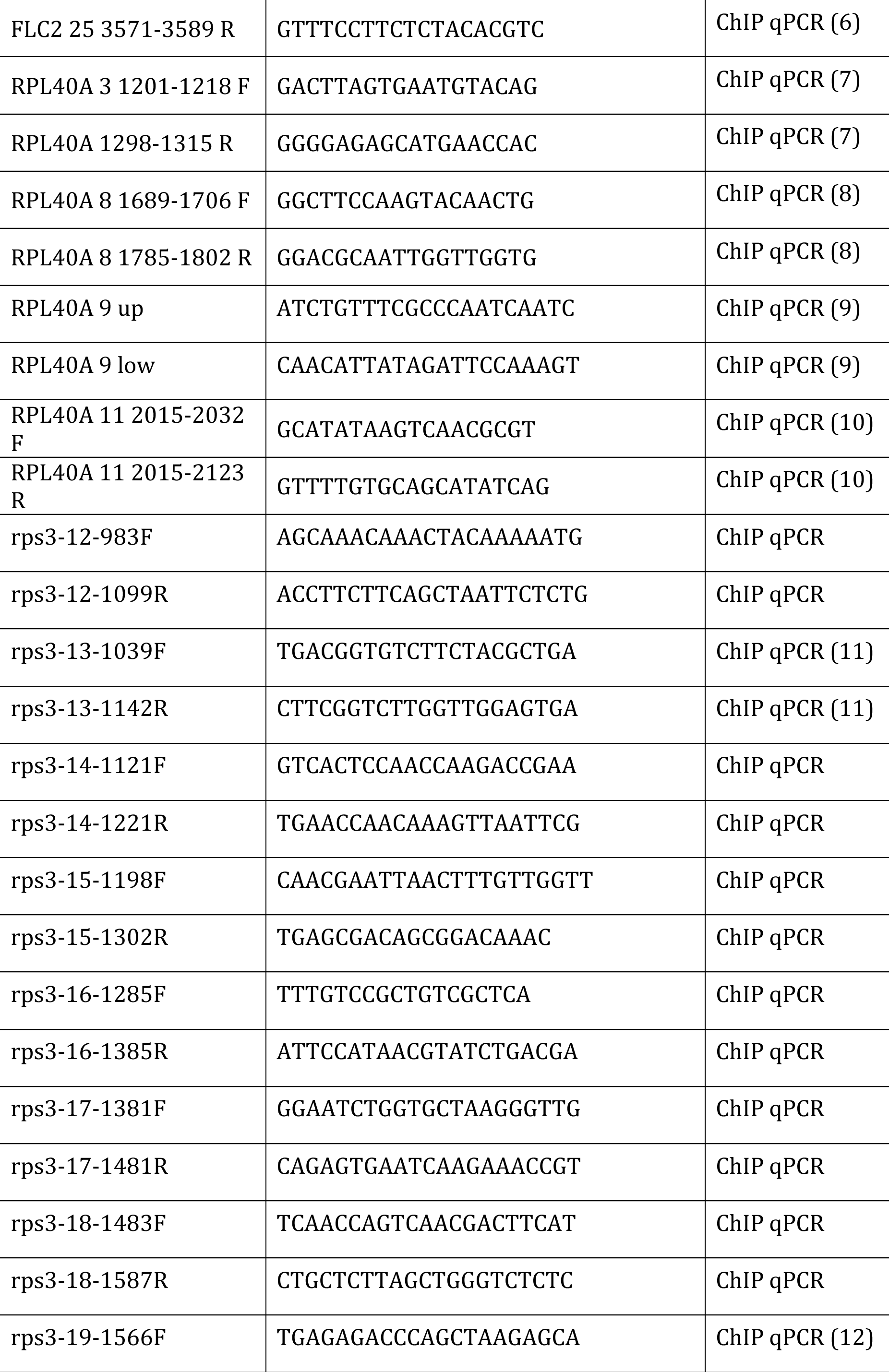
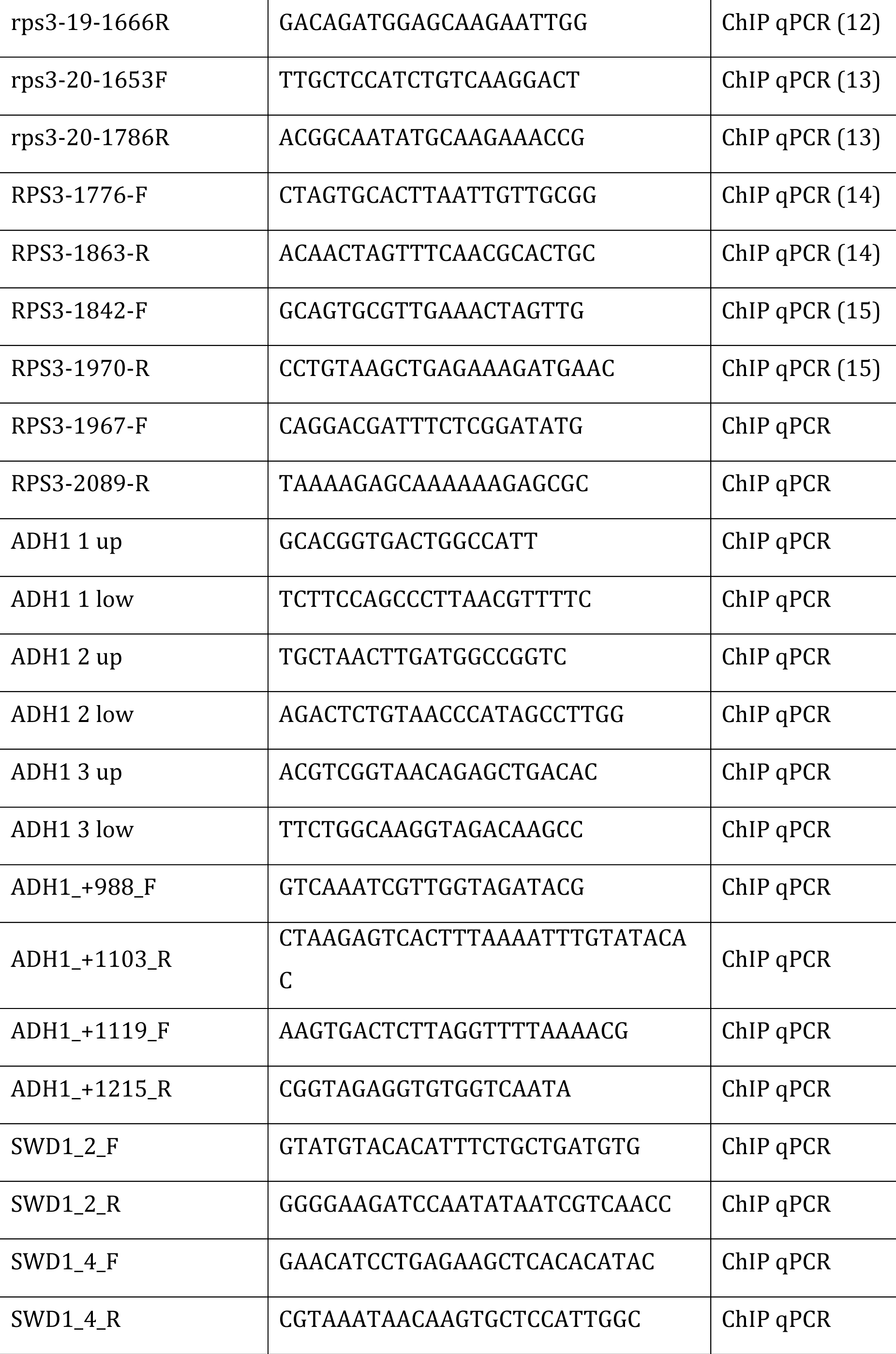
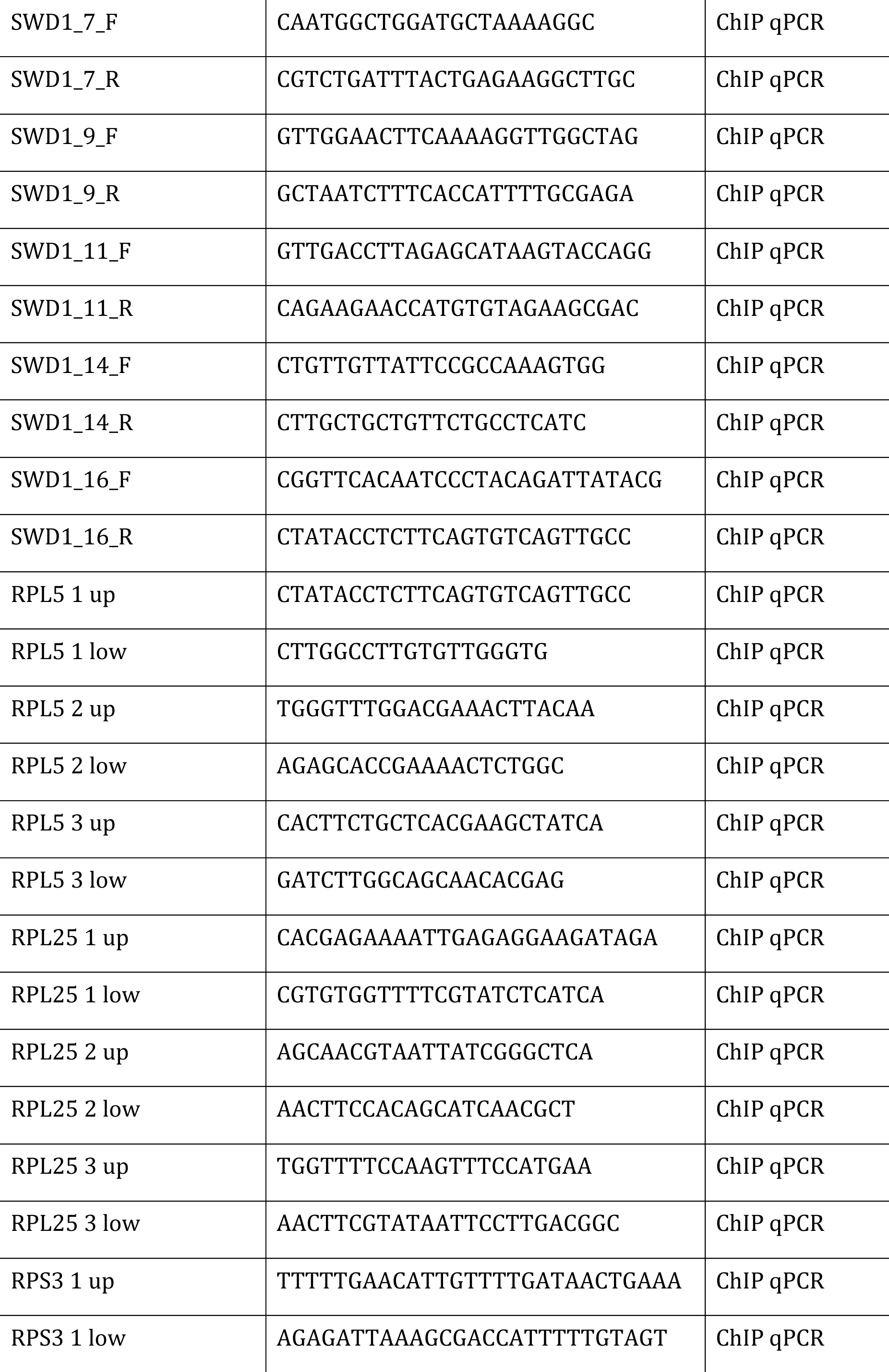
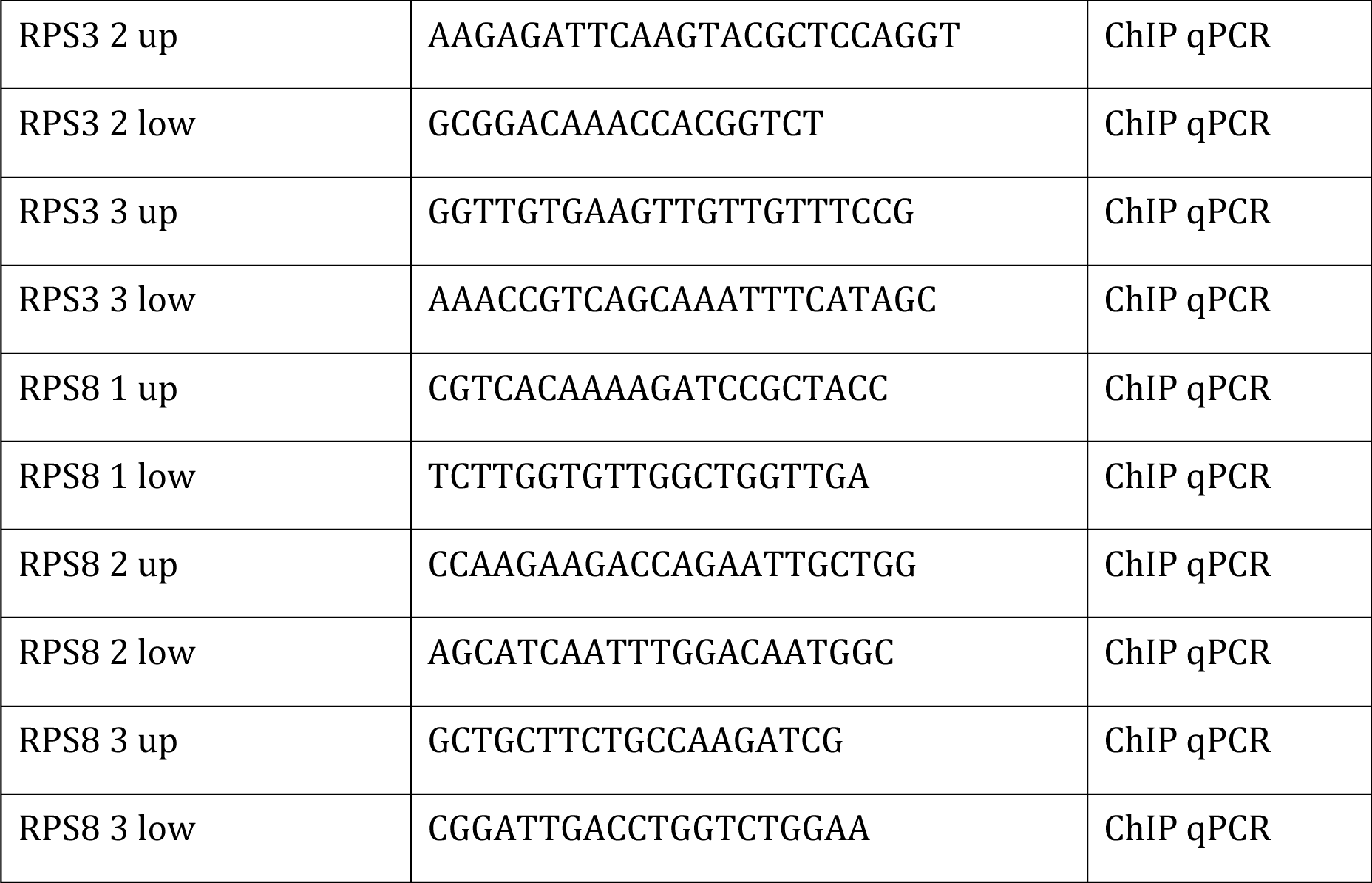
Primers used in this work.

